# Establishment of CRISPR/Cas9-based knock-in in a hemimetabolous insect: targeted gene tagging in the cricket *Gryllus bimaculatus*

**DOI:** 10.1101/2021.05.10.441399

**Authors:** Yuji Matsuoka, Taro Nakamura, Takahito Watanabe, Austen A. Barnett, Sayuri Tomonari, Guillem Ylla, Carrie A. Whittle, Sumihare Noji, Taro Mito, Cassandra G. Extavour

## Abstract

Studies of traditional model organisms like the fruit fly *Drosophila melanogaster* have contributed immensely to our understanding of the genetic basis of developmental processes. However, the generalizability of these findings cannot be confirmed without functional genetic analyses in additional organisms. Direct genome editing using targeted nucleases has the potential to transform hitherto poorly-understood organisms into viable laboratory organisms for functional genetic study. To this end, here we present a method to induce targeted genome knock-out and knock-in of desired sequences in an insect that serves as an informative contrast to *Drosophila*, the cricket *Gryllus bimaculatus*. The efficiency of germ line transmission of induced mutations is comparable to that reported for other well-studied laboratory organisms, and knock-ins targeting introns yield viable, fertile animals in which knock-in events are directly detectable by visualization of a fluorescent marker in the expression pattern of the targeted gene. Combined with the recently assembled and annotated genome of this cricket, this knock-in/knock-out method increases the viability of *G. bimaculatus* as a tractable system for functional genetics in a basally branching insect.

## Introduction

In what is often called the “post-genomic era,” (Wainberg et al., 2021), massive advances in nucleic acid sequencing chemistry over the last two decades have given scientists access to greater volumes of gene sequence data than ever before (Kulski, 2016; Papageorgiou et al., 2018). However, this wealth of genomic information has highlighted two major gaps in our understanding of gene function and evolution. First, comparative genomic data and increased taxon sampling in functional genetic, developmental and cellular biology have revealed that the biology of many traditional laboratory model organisms is not representative of the broader clades to which they belong (Goldstein and King, 2016). Second, our ability to accurately deduce gene function from sequence data is limited to those genes that display high sequence and structural conservation (Ashburner et al., 2000; Consortium et al., 2020), and tools for manipulating gene function have been developed for only a small fraction of organisms (Russell et al., 2017). Addressing these problems calls for both increased taxon sampling and development of techniques to enable targeted alteration of gene function in understudied organisms. Here we address both of these issues by developing a method for targeted genome editing, including both knock-out and knock-in editing, in a basally branching insect model organism, the cricket *Gryllus bimaculatus*.

*G. bimaculatus* is an emerging model organism in a variety of fields of biology (Horch et al., 2017a; Kulkarni and Extavour, 2019). Ease of husbandry (Horch et al., 2017b), detailed developmental staging tables (Donoughe and Extavour, 2016), established gene expression analysis methods (Horch et al., 2017b), and an assembled and annotated genome (Ylla et al., 2021) make this cricket a highly amenable hemimetabolous laboratory model system (Kulkarni and Extavour, 2019). In contrast to the ontogenetically derived model *Drosophila melanogaster*, many aspects of cricket embryogenesis are thought to resemble putative ancestral developmental modes of insects (Davis and Patel, 2002). For example, the function of several axial patterning genes has been analyzed and compared to that of their *D. melanogaster* homologues, revealing that the gene regulatory networks governing axial patterning have undergone considerable evolutionary change across insects (Mito et al., 2007; Mito et al., 2008; Matsuoka et al., 2015).

*G. bimaculatus* is also used for the analysis of gene function in tissue regeneration (Mito et al., 2002; Nakamura et al., 2007; Bando et al., 2013), as this cricket can regenerate amputated organs, including legs and antennae, after several rounds of juvenile molts. In addition, *G. bimaculatus* is used for the analysis of gene function and neuronal circuits in neuronal activity, including learning, memory and circadian clocks (Hedwig and Sarmiento-Ponce, 2017; Matsumoto et al., 2018; Mizunami and Matsumoto, 2017; Tomiyama et al., 2020). Moreover, several species of cricket are being farmed as a new food source for humans because of their high protein and nutrient content (Huis et al., 2013).

Genome editing techniques using artificial nucleases were previously established in *G. bimaculatus* (Watanabe et al., 2012). However, construction of artificial nucleases is laborious. More recently, use of the <u>c</u>lustered regulatory interspaced <u>s</u>hort <u>p</u>alindromic repeat (CRISPR)/associated Cas9 nuclease (CRISPR/Cas9) has emerged and been verified as an efficient tool for genome editing in several arthropod species, including the fruit fly *Drosophila melanogaster* (Gratz et al., 2013), the beetle *Tribolium castaneum* (Gilles et al., 2015), the mosquito *Aedes aegypti* (Kistler et al., 2015), multiple butterfly species (Li et al., 2015; Matsuoka and Monteiro, 2018; Zhang et al., 2017), the amphipod *Parhyale hawaiensis*, (Martin et al., 2016), the clonal raider ant *Ooceraea biroi* (Trible et al., 2017), the European honeybee *Apis mellifera* (Kohno and Kubo, 2018), and the milkweed bug *Oncopeltus fasciatus* (Reding and Pick, 2020). Among crickets, we and others have reported CRISPR/Cas9-mediated knockouts in *Acheta domesticus* (Dossey e tal. 2023) and in *G. bimaculatus* (Bai et al. 2023, Nakamura et al. 2022). Knock0in has also been reported for *A. domesticus* (Dossey et al. 2023), but this approach has not been used to generate the types of enhancer trap or protein trap lines that have proven so beneficial to advancing developmental biology research in other insects (Morin et al. 2001). Here we briefly review this technique, investigate and demonstrate its utility for targeted gene trap knock-in genome modification in *G. bimaculatus*, the most widely used orthopteran model for developmental biology.

In the CRISPR/Cas9 system, short guide RNAs (sgRNAs) recruit Cas9 nuclease to the target sequence, and Cas9 then introduces a double strand break (DSB) at the target sequence. The presence of DSBs triggers the activity of the cell’s DNA repair machinery, either <u>n</u>on-<u>h</u>omologous <u>e</u>nd joining (NHEJ) or <u>h</u>omology <u>d</u>irected repair (HDR). NHEJ is an error-prone machinery, such that insertions or deletions can be generated at the break point (Branzei and Foiani, 2008). By using artificial nucleases to trigger NHEJ, we previously succeeded in generating *G. bimaculatus* mutant lines (Watanabe et al., 2012). HDR, however, would offer more precise repair machinery, as the break is repaired through use of a homologous template. By supplying a donor template containing sequence homologous to the target, in principle a desired donor sequence can be integrated into the genome though HDR. Such gene knock-ins, while highly desirable for detailed analysis of the function of genomic regions, are more difficult to achieve than gene knock-outs because of the low efficiency of HDR in eukaryotes (Hagmann et al., 1998). Although success with HDR has been reported in some insects including the silk moth *Bombyx mori* (Ma et al., 2014; Zhu et al., 2015), multiple mosquito species (Hammond et al., 2016; Kistler et al., 2015; Purusothaman et al., 2021) and mosquito cell lines (Rozen-Gagnon et al., 2021), the beetle *Tribolium castaneum* (Gilles et al., 2015), the medfly *Ceratitis capitata* (Aumann et al., 2018), and the squinting bush brown butterfly *Bicyclus anynana* (Connahs et al., 2022), our attempts at HDR-based knock-in techniques have never succeeded in *G. bimaculatus* (unpublished observations). Recently, an efficient gene knock-in method through NHEJ was developed in the zebrafish *Danio rerio* (Auer et al., 2014). In this method, both the genome and the donor vector are cleaved *in vivo*, then the terminal genomic and donor sequences are combined through NHEJ. The method is efficient and can integrate longer constructs into the genome than knock-ins achieved through HDR (Auer et al., 2014, Yoshimi et al., 2016). Bosch and colleagues (Bosch et al., 2019) subsequently reported that this knock-in strategy also works in *D. melanogaster*.

Here, we present evidence that the CRISPR/Cas9 system functions efficiently in *G. bimaculatus*. We demonstrate the utility of this technique for developmental biology by performing functional analysis of the *G. bimaculatus* orthologues of the Hox genes *Ultrabithorax* (*Gb-Ubx*) and *abdominal-A* (*Gb-abd-A*). Furthermore, using a donor vector containing an autonomous expression cassette, we demonstrate that gene knock-in by a homology-independent method works efficiently in *G. bimaculatus*. We show that this homology-independent gene knock-in method can be applied to identify mutant individuals simply by detecting marker gene expression in this cricket. Efficient targeted genome editing, now including both knock-out and gene-tagging knock-in techniques, will pave the way for making this cricket a much more sophisticated model animal for functional genetic laboratory studies.

## Results

### Targeted mutagenesis of the Gb-lac2 locus

To determine whether the CRISPR/Cas9 system was functional in the cricket, we first tried to perform a targeted gene knock-out of the *laccase 2* (*Gb*-*lac2*) gene (Table 1), which regulates tanning of the arthropod cuticle following molting (Arakane et al., 2005). We chose this gene because of its easily detectable loss-of-function phenotype, and because we had previously successfully generated stable mutant lines for this gene by using artificial nucleases (Watanabe et al., 2012). sgRNA target sites were first chosen using the CasOT tool (Xiao et al., 2014), and then we chose the final target sequence from among these candidate sequences based on the number of mismatches relative to the other sequences in the genome (> 3 mismatches in the whole sgRNA sequence) and GC content (70 ± 10% for the whole sgRNA sequence). Based on these criteria, we designed sgRNAs against the fifth exon of *Gb*-*lac2*, which is close to the target regions of the previously reported artificial nuclease experiment (Fig. 1D; Table 2) (Watanabe et al., 2012).

**Table 1.** *G. bimaculatus* genes in this study disrupted by targeted genome modification

| *Gene* | *Functions* | *Tissue distribution* | *Phenotype of genome-modified knockout/knockdown cricket* | *Refs* |
| --- | --- | --- | --- | --- |
| *Laccase 2* | Phenol oxidase, cuticle tanning (sclerotization and pigmentation) | No data | *Lac2* knock-out nymphs show defect in pigmentation. | (Watanabe et al., 2014); this study |
| *Ultrabithorax* | Enlargement of T3 leg<br>Identity of A1 pleuropodia | T3 and A1 segment | - *Ubx* knock-out embryos show partial transformation of A1 pleuropodia into T3 thoracic leg, and of T3 thoracic leg into T2 thoracic leg.<br>- Embryonic lethal. | This study |
| *abdominal-A* | Repression of leg formation in abdomen | Abdomen | - *abd-A* knock-out embryos show generation of leg-like structures on the abdomen.<br>- *abd-A* knock-out nymphs show fusion of abdominal segments.<br>- *abd-A* knock-out female adults generated ectopic ovipositors and had defects in oviducts and uterus attachment. | This study |

**Figure 1.**
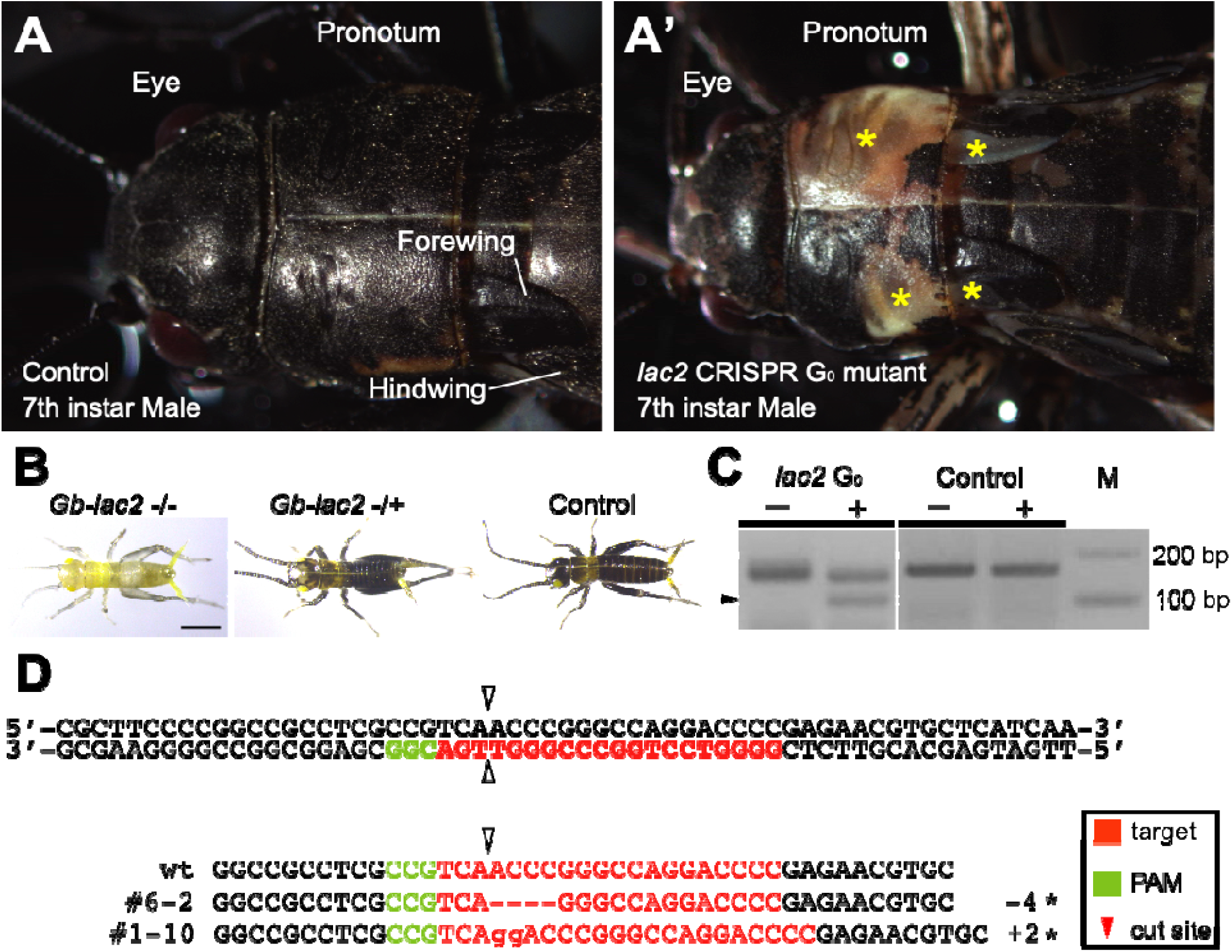
*Gb-laccase2* knock-out G_0_ and G_1_ phenotypes. (A) The cuticle of adult wild type (WT) *G. bimaculatus* is uniformly dark brown or black. Head and anterior thorax shown in dorsal view. (A’) *Gb-lac2* gene somatic mutagenesis in G_0_ animals can be detected by the presence of white or light brown spots of cuticle (asterisks). (B) Representative cuticle phenotypes of G_1_ *Gb-lac2* mutant nymphs at one day after hatching. Homozygous mutants (-/-) showed homogeneous pale brown cuticle. Heterozyous mutants (-/+) showed slight reduction in black pigmentation. Control first instar nymphs have dark melanized cuticle by one day after hatching. Scale bar:1 mm. (C) Surveyor^TM^ assay result. The control experiment, in which the PCR product was amplified from the genome of a wild type control individual, did not produce any band of the expected size after Surveyor^TM^ endonuclease treatment. In contrast, the PCR product amplified from the genome of CRISPR reagent-injected animals included small fragments of the expected size after Surveyor^TM^ endonuclease treatment (black arrowhead). M: electrophoresis marker. (D) Sequence analysis of *Gb-lac2* mutant alleles #6-2 and #1-10 induced by the CRISPR/Cas9 system. Top row: wild-type sequences; green: Protospacer Adjacent Motif (PAM) sequence; red: target sequence; arrowheads: predicted double strand break site. Asterisks at right show induced frame-shift mutations.

**Table 2.**
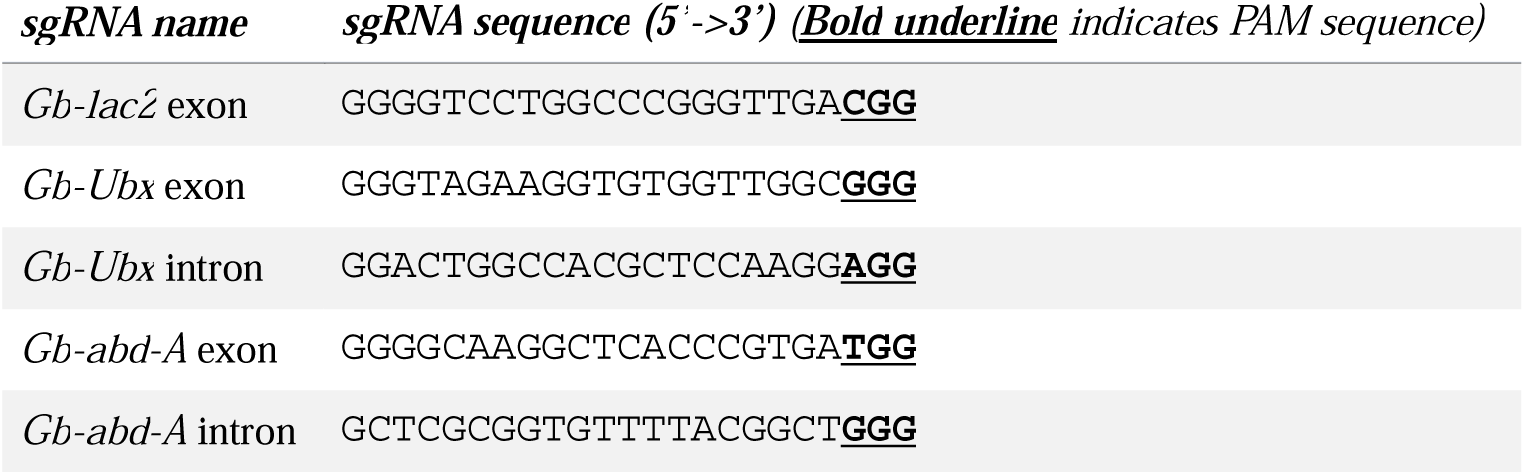
sgRNA sequences used in this study

We co-injected 0.5 μg/μl sgRNA and 0.5 μg/μl of Cas9 mRNA into 232 fertilized cricket eggs within 1-3 hours after egg laying (h AEL) (Table 3). Five days after injection, we evaluated the frequency of mutant alleles in individual eggs using the Surveyor^TM^ nuclease assay (Qiu et al., 2004) (see Materials and Methods for detailed mechanism and procedure). We detected cleaved fragments in 13 out of 29 eggs examined (Table 3, Fig. 1C).

**Table 3.** Efficiency of CRISPR/Cas9-mediated genome editing in *G. bimaculatus. nd = no data*

|  | *Shown in Figure* | *# eggs injected* | *# injected embryos with eGFP expression or phenotype* | *# embryos developed by 7d AEL (% of injected embryos)* | *# hatched nymphs (% of embryos developed by 7d AEL)* | *# fertile adults (% of nymphs hatched)* | *% injected embryos yielding fertile adults* | *# fertile adults showing germ line transmission* | *% fertile adults showing germ line transmission* | *% injected embryos yielding fertile adults showing germline transmission* |
| --- | --- | --- | --- | --- | --- | --- | --- | --- | --- | --- |
| *Gb-Lac2*<sup>CRISPR</sup> KO | 1 | 232 | nd | nd | nd | 38 (nd) | 16.4% | 18 | 47.4% | 7.8% |
| *Gb-Ubx*<sup>CRISPR</sup> KO | 2, S1, S2 | 167 | nd | nd | nd | 10 (nd) | 59.9% | 6 | 60.0% | 3.6% |
| *Gb-Ubx*<sup>KI-exon</sup> | 3, S3, S4 | 85 | 4 (4.7%) | 58 (68.2%) | 30 (51.7%) | 25 (83.3%) | 29.4% | 1 | 4.0% | 1.2% |
| *Gb-abd-A*<sup>KI-exon</sup> 1<sup>st</sup> trial | 4, S2, S3, S4, S5, S6 | 47 | 5 (10.6%) | 38 (80.8%) | 9 (23.7%) | 4 (44.4%) | 8.5% | 1 | 25.0% | 2.1% |
| *Gb-abd-A*<sup>KI</sup> exon 2<sup>nd</sup> trial | n/a | 41 | 0 (0.0%) | 36 (87.8%) | 22 (61.1%) | 2 (9.1%) | 4.9% | 1 | 50.0% | 2.4% |
| *Gb-abd-A*<sup>KI-intron</sup> | 5, S5, S6 | 100 | 2 (2.0%) | 77 (77.0%) | 47 (61.0%) | 28 (59.6%) | 28.0% | 1 | 3.6% | 1.0% |
| *Gb-Ubx*<sup>KI-intron</sup> | S7 | 73 | 5 (6.8%) | 62 (84.9%) | 59 (95.2%) | 22 (37.3%) | 30.1% | 2 | 9.1% | 2.7% |
| Control (Donoughe and Extavour, 2016) | n/a | 42 | n/a | 40 (95.2%) | 28 (70.0%) | nd | nd | n/a | n/a | n/a |
| Control (Ewen-Campen et al., 2012) | n/a | 78 | n/a | nd | 64 (82.0%) | nd | nd | n/a | n/a | n/a |

We observed mosaic pigmentation of the cuticle in 92% of G_0_ hatchlings that emerged from the individual eggs injected with the *Gb*-*lac2* sgRNA, consistent with Cas9-mediated interruption of the *Gb*-*lac2* gene in some, but not all, somatic cells of the G_0_ hatchlings. We raised these hatchlings to adulthood (Fig. 1A’), and crossed these G_0_ adults with wild type crickets of the opposite sex, to determine the efficiency of germ line transmission of the Cas9-induced *Gb*-*lac2* mutations to the G_1_ generation. We found that 47.4% of those injected with the *Gb*-*lac2* gRNA transmitted the mutation to their offspring (Fig. 1B). To determine the nature of the *Gb*-*lac2* Cas9-induced alleles, we isolated genomic DNA from each line and analyzed the sequence of the *Gb*-*lac2* locus. We found that several different types of indel mutations were introduced at the target locus (Fig. 1D). These results indicate that this CRISPR/Cas9-mediated genome editing system is functional in the cricket.

### Targeted mutagenesis of the Gb-Ubx locus via knock-out

To compare phenotypes obtained with targeted gene disruption to those obtained with the RNA interference (RNAi) method that has hitherto been the most common method of performing functional genetics in this cricket (Mito and Noji, 2008), we used the CRISPR/Cas9 system to perform functional analyses of the *G. bimaculatus* ortholog of the Hox gene *Ultrabithorax* (*Gb*-*Ubx*) (Table 1). We previously examined RNAi-induced phenotypes for *Gb-Ubx* in developing abdominal segments (Barnett et al., 2019), providing a basis for comparison with CRISPR-induced mutants. We designed sgRNA for a sequence within an exon upstream of the homeodomain (Fig. 2A), and co-injected 0.5 μg/μl of this sgRNA and 1 μg/μl Cas9 mRNA into 167 fertilized cricket eggs within 1-3 h AEL. Seven days following injection, we extracted genomic DNA from a small subset of injected eggs and performed the Surveyor^TM^ assay to determine the efficiency of gene targeting. We found that *Gb-Ubx* mutations had been induced in all examined eggs (n=16) (Fig. 2B). The remaining 151 injected G_0_ embryos gave rise to ten adults, which we backcrossed individually to wild type adults of the opposite sex. We randomly chose approximately 30 G_1_ eggs from each of the ten G_0_ crosses, extracted genomic DNA from the pooled embryos, and performed the Surveyor^TM^ assay. We found that six out of ten G_0_ crickets transmitted *Gb-Ubx* mutations to the next generation. We selected one of the six G_1_ *Gb-Ubx^CRISPR^* lines, which had a frame-shift mutation in the *Gb-Ubx* locus, for further phenotypic analysis. These *Gb-Ubx^CRISPR^* mutants displayed two different classes of phenotype: (1) *Contraction of the T3 leg*. Wild type *G. bimaculatus* adults have large, conspicuous T3 jumping legs. However, heterozygous *Gb-Ubx^CRISPR^* mutants had smaller T3 legs than wild type animals (Fig. 2C). Homozygous *Gb-Ubx^CRISPR^* mutants obtained in the G_2_ generation had T3 legs that were even smaller than those of heterozygotes, almost the same size as T1/T2 legs (Fig. 2C). This specific phenotype is not directly comparable with previously reported *Gb-Ubx* RNAi experiments (Barnett et al., 2019), because embryos in those experiments were not reared to hatching. However, the *Gb-Ubx^CRISPR^* phenotypes were in good correspondence with those previously observed for *Ubx* RNAi in the cricket *Acheta domesticus* (Mahfooz et al., 2007). (2) *Transformation of the A1 appendage*. Wild type *G. bimaculatus* germ band stage embryos possess two appendage-like organs on the A1 segment called pleuropodia (Fig. 2D; Rathke, 1844; Wheeler, 1892). Instead of the pleuropodia present in wild type adults, the appendage outgrowths on the A1 segment of homozygous *Gb-Ubx^CRISPR^* mutants were transformed towards leg-like structures, which is consistent with detection of expression of *Gb-Dll* and other leg patterning genes in the leg-like structures. (Fig. 2D and Supplemental Figure 1). This phenotype matches that previously observed in *Gb-Ubx* RNAi embryos (Fig. 2E; Barnett et al., 2019). *Gb-Ubx^CRISPR^* heterozygous mutants were fertile but homozygous mutants were lethal. Therefore, to maintain this line, *Gb-Ubx^CRISPR^*heterozygous mutant animals of separate sexes were crossed to each other, and we performed the Surveyor^TM^ assay to isolate heterozygous mutants from among their offspring.

**Figure 2.**
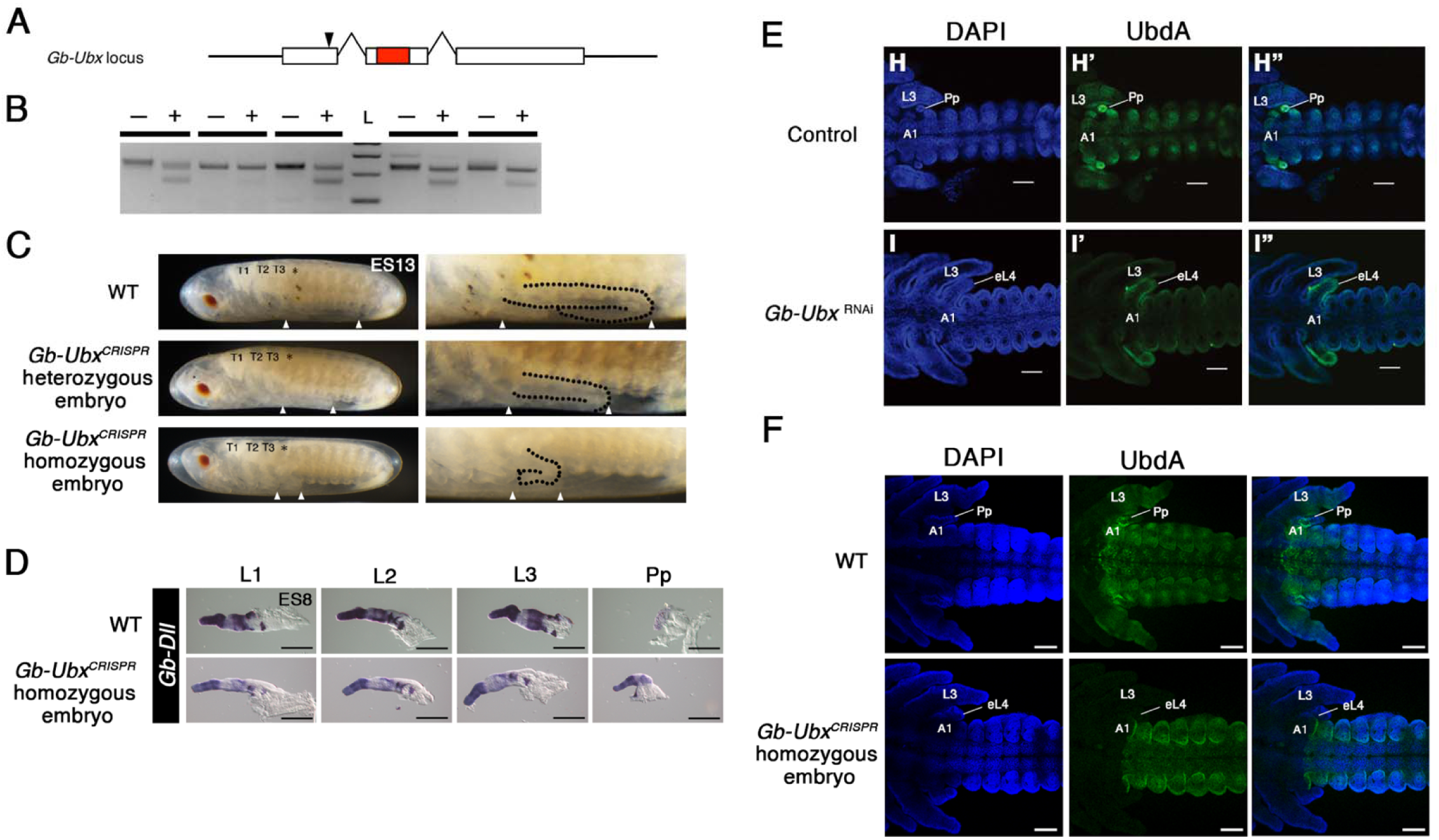
Knock-out vs knock-down phenotype of *Gb-Ubx*. (A) Schematic diagram of *Gb-Ubx* locus. White boxes: exons; red box: homeodomain; black arrowhead: sgRNA target site. (B) Surveyor^TM^ assay results with G_0_ eggs. Plus (+) indicates the PCR products digested by Surveyor^TM^ nuclease; minus (-) indicates the PCR products with no digestion (no nuclease added). L = ladder. (C) Phenotype of heterozygous and homozygous *Gb-Ubx^CRISPR^* mutant stage 11 embryos. Anterior is to the left here and in all other figures. The size of the T3 leg (region between white arrowheads) was decreased mildly and severely in heterozygous and homozygous mutants, respectively. T3 legs are delineated with black dotted line in higher magnification images at right figures. Anterior (leftmost) white arrowhead marks the posterior end of T3 segment. Posterior (rightmost) white arrowhead marks the junction of femur and tibia. (D) Expression pattern of *Gb-Dll* in thoracic and abdominal appendages of wild type and homozygous *Gb-Ubx^CRISPR^* mutant embryos. Tarsus is to the left (distal); tibia is to the right (proximal). In wild type embryos, *Gb*-*Dll* is expressed strongly in the presumptive tarsus of developing T1, T2 and T3 legs (L1, L2 and L3 respectively). In the tibia of T3, different from T1 and T2, a sharply defined border between a distal domain of high expression is detectable. In homozygous *Gb-Ubx^CRISPR^*mutant embryos, this T3-specific expression pattern was not detected. In wild type embryos, *Gb-Dll* is ubiquitously expressed in the pleuropodia, but it showed the leg-like expression pattern in homozygous *Gb-Ubx^CRISPR^* mutant embryos. (E) Ubx/Abd-A (UbdA) protein expression pattern in *Gb-Ubx^RNAi^* stage 8 embryos. In stage 8 *Gb-Ubx^RNAi^*embryos, UbdA protein expression was undetectable in the T3 leg (L3) but was still detected in the A1 segment. (F) Ubx/Abd-A (UbdA) protein expression pattern in homozygous *Gb-Ubx^CRISPR^* stage 9 embryos. In *Gb-Ubx*^CRISPR^ embryos, only the T3/A1 Gb-Ubx expression domain was undetectable. Scale bar: 100 μm. Pp: pleuropodium. Embryonic staging as per (Donoughe and Extavour, 2016).

To confirm whether the production of Gb-Ubx protein was indeed disrupted by these CRISPR-induced mutations, we performed immunostaining with the “UbdA” monoclonal antibody FP6.87 (Kelsh et al., 1994), which recognizes both Ubx and Abd-A proteins, and was previously reported to cross-react in multiple *Gryllus* species including *G. bimaculatus* (Barnett et al., 2019; Mahfooz et al., 2004). In wild type embryos, the UbdA antibody revealed the expected combination of the Gb-Ubx and Gb-Abd-A expression patterns (Fig. 2E and 2F; Barnett et al., 2019; Mahfooz et al., 2004). In *Gb-Ubx^CRISPR^* embryos, however, while the UbdA expression domain was retained in the abdomen where *Gb-abd-A* is expressed, expression was undetectable in the T3 and A1 segment where *Gb-Ubx* is expressed (Fig. 2F), suggesting that the CRISPR-induced *Gb-Ubx* mutations interfered with Gb-Ubx protein production. We note that in previous reports, UbdA expression was not affected in *Gb-Ubx^RNAi^* embryos, which might be because RNAi does not disrupt target gene expression completely (REF).

We further characterized the phenotype of *Gb-Ubx^CRISPR^*embryos by observing the *Gb*-*Dll* expression pattern (Fig. 2D). In wild type embryos, *Gb*-*Dll* is expressed strongly in the presumptive tarsus of all three developing T1, T2 and T3 legs (Fig. 2D). In the tibia, however, *Gb-Dll* is expressed differently in T1/T2 and T3 legs. Specifically, in T3 there is a sharply defined border between a distal domain of high expression and a proximal domain of lower expression, but T1 and T2 lack such a border and display moderate proximal expression as well as strong distal expression in (Niwa et al., 1997) (Fig. 2D). In *Gb-Ubx^CRISPR^* embryos, *Gb-Dll* expression in the presumptive tibia of the T3 leg lacked a strong boundary between high and low tibial expression levels, and instead resembled the expression pattern of wild type T1 or T2 legs (Fig. 2D). In addition, the T3 leg-specific pattern of multiple leg-patterning genes was absent in *Gb-Ubx^CRISPR^*embryos, which is consistent with disruption of the *Gb*-*Ubx* locus (Supplementary Fig.1). Taken together, these results suggest that the CRISPR/Cas9 system induced mutations specifically at the *Gb-Ubx* locus, which disrupted *Gb-Ubx* function.

### In-depth analysis of mutagenesis profile for the CRISPR/Cas9 system in G. bimaculatus

To optimize the genome editing procedure, we wished to evaluate whether and how the timing of injection affected NHEJ mutagenesis. For detailed assessment of this mutagenesis, we therefore performed in-depth analysis of the CRISPR mutants using next generation sequencing.

Our previous study had revealed early cellular dynamics during cricket embryogenesis (Nakamura et al., 2010), allowing us to assess whether specific mutagenesis events were correlated with cellular behaviors during early development. As in *D. melanogaster* (Foe and Alberts, 1983) early mitotic divisions in *G. bimaculatus* embryos are syncytial, meaning that mitosis takes place without cytokinesis, resulting in multiple energids (nuclei surrounded by aqueous cytoplasm but lacking a unique lipid bilayer) within a single cell membrane (Donoughe et al. 2021; Donoughe and Extavour, 2016; Nakamura et al., 2010; Sarashina et al., 2003). To evaluate whether and how the timing of injection affected mutagenesis outcomes, we chose four early embryonic time points following the one-hour (1h) embryo collection period, as follows (Supplementary Fig. 2A): (1) At the 1h injection time point, energids start to migrate from the center of the egg to the cortex. (2) At the 3h injection time point, energids continue to become distributed throughout the yolk, accompanied by mitotic cycles. (3) At the 5h injection time point, energids have become nearly uniformly distributed throughout the egg cortex and begin tangentially oriented nuclear division. (4) The 9h injection time point is one to eight hours before cellularization (Donoughe and Extavour, 2016). We co-injected 0.5 μg/μl of the *Gb-Ubx* sgRNA, the *Gb-lac2* sgRNA or an sgRNA targeting *abdominal-A* (*Gb-abd-A*); see “*Knock-in of donor vector sequence at the Gb-abd-A locus*” below) described above, and 1 μg/μl of Cas9 mRNA into the eggs at each of these time points. At 5d AEL, we isolated genomic DNA from three individual embryos for each injection time point and used it for amplicon sequencing. We examined the sequences of the on-target site and of the single highest predicted potential off-target site for each of the *Gb-Ubx* and *Gb-abd-A* genes. For each sample we performed amplicon sequencing with three replicates.

For *Gb-Ubx* on-target gene disruptions, we found that the rate of NHEJ-induced mutations decreased with the age of the embryo at injection (Supplementary Fig. 2B). The rate of NHEJ-induced mutations at the studied off-target site was less than 1.3% for all injection time points (Supplementary Fig. 2C), suggesting that off-target effects may be minimal in this system. The same trend was also observed for *Gb-abd-A* on-target mutations (Supplementary Fig. 2D-E; see section “*Targeted mutagenesis of the Gb-abd-A locus via knock-in”* below). This result is well correlated with the phenotypic severity observed in the G_0_ hatchlings emerging from the *Gb-lac2^CRISPR^* embryos. *Gb-lac2^CRISPR^*embryos injected at the two earlier time points (1h and 3h) gave rise to hatchlings with broad patches of white cuticle (Supplementary Fig. 2H). In contrast, the embryos injected at 5h showed milder phenotypes (Supplementary Fig. 2H) and the embryos injected at 9h showed only little detectable phenotype (Supplementary Fig. 2H).

### Knock-in of donor vector sequence at the Gb-Ubx locus

In addition to targeted sequence deletions, targeted sequence knock-in is a highly desirable technique that would expand our ability to understand the functions of genomic regions of interest. We had previously attempted to achieve targeted gene knock-ins through homology-dependent repair, but this method has not worked in *G. bimaculatus* in our hands to date (data not shown). In a homology-independent knock-in method reported for *D. rerio* and *D. melanogaster* (Auer et al., 2014; Bosch et al., 2019), both genome and donor vector are cleaved *in vivo*, then the cut ends of genome and donor vector are combined through NHEJ. This method is more efficient than the homology-dependent method, potentially because NHEJ is highly active throughout the cell cycle in eukaryotes (Hagmann et al., 1998). However, due to the nature of NHEJ, the orientation of integration of the donor vector sequence cannot be controlled. In addition, indel mutations are generated at the junction point. To try to circumvent these issues, which might otherwise prevent functional knock-in, we generated a donor vector containing an autonomous expression cassette comprising the *G. bimaculatus actin* (*Gb-act*) promoter followed by the *eGFP* coding sequence (Nakamura et al., 2010). As a sgRNA recognition site, we included a partial *DsRed* gene sequence (Auer et al., 2014), which is native to the coral *Discosoma sp.* (Baird et al., 2000) and not present in the cricket genome. We predicted that successful knock-in of this donor sequence into the genome would result in eGFP expression being driven by the *Gb-act* promoter regardless of the insert’s orientation or any potential induced indel mutations. To try to further increase the utility of this tool to facilitate identification of targeted gene disruptions, we targeted knock-in of the donor sequence to an exon of the target gene, which we anticipated would result in disruption of target gene function. Our goal was to be able to identify such successfully knocked-in individuals by detectable eGFP expression in the known expression domains of the target gene.

We chose the *Gb-Ubx* locus for this targeted knock-in strategy and used the same sgRNA as that used for the knock-out experiment described above (see section “*Targeted mutagenesis of the Gb-Ubx locus via knock-out*”, Fig. 2A). We co-injected 50 ng/μl of sgRNA for the *Ubx* locus, 50 ng/μl of sgRNA for the donor vector, 100 ng/μl of Cas9 mRNA, and 100 ng/μl of donor vector into fertile cricket eggs. By seven days after injection, four out of 85 injected embryos (4.7%) showed mosaic eGFP expression in the T3 trunk and leg (Fig. 3B’). Of the 85 injected embryos, 18 individuals (21.2%) grew to adulthood. We crossed them individually with wild type counterparts of the opposite sex and evaluated eGFP expression in their offspring. One out of the 18 G_0_ crickets (5.6%) produced G_1_ embryos with eGFP expression in a pattern identical to that of *Gb-Ubx* (Fig. 3C’; Barnett et al., 2019; Matsuoka et al., 2015; Zhang et al., 2005). The eGFP expression was detectable through the eggshell even at late embryonic stages (Fig. 3C’). At adult stages, G_1_ knock-in crickets showed detectable eGFP expression in the hind wing and T3 legs (Supplementary Fig. 3A’ and B’).

**Figure 3.**
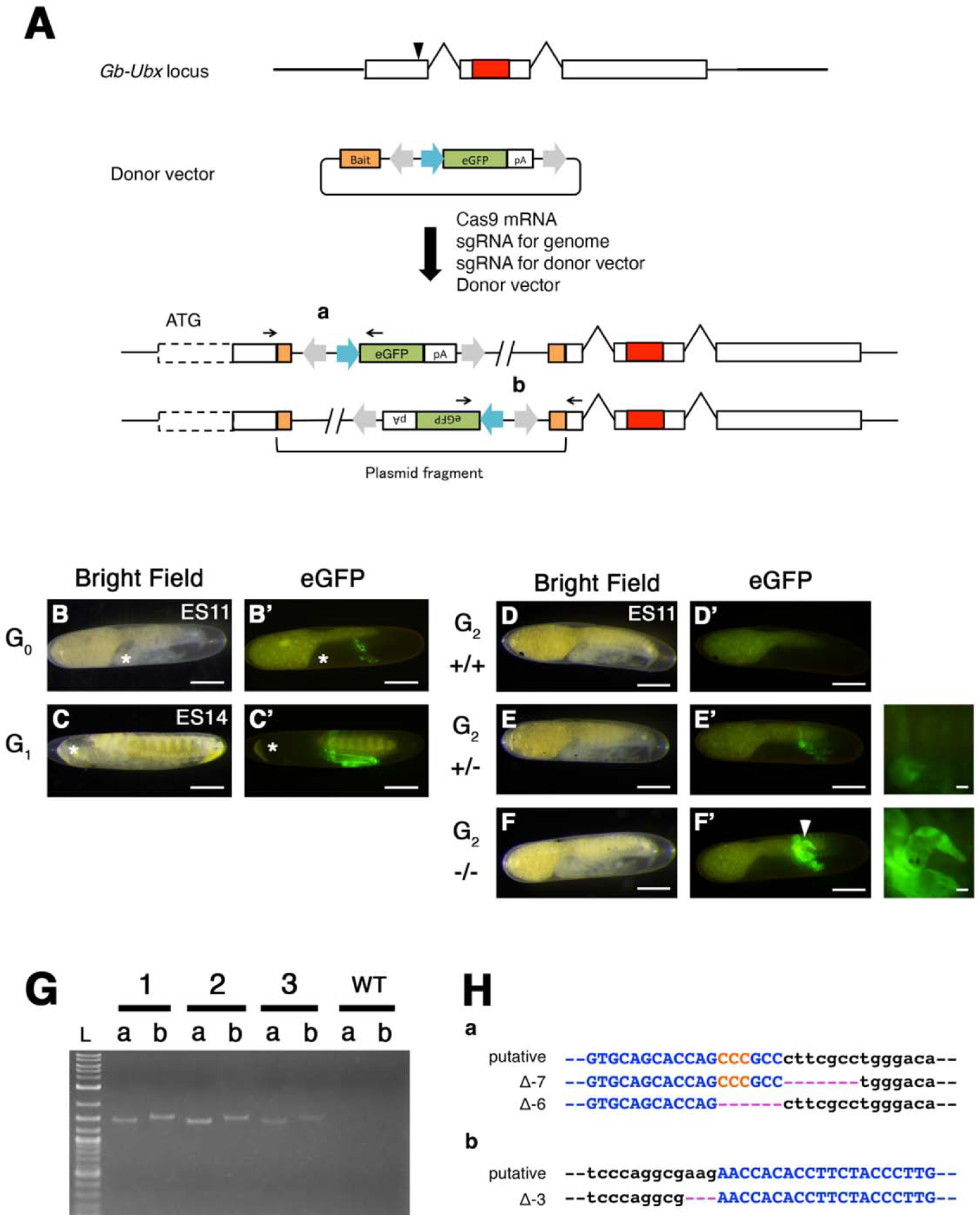
Exonic knock-in/Knock-out against *Gb-Ubx*. (A) Scheme of knock-in experiment targeting a *Gb-Ubx* exon. White boxes: exons; red box: homeodomain; black arrowhead: sgRNA target site. Donor vector contains bait sequence (orange box) and expression cassette with the following elements: *Gryllus actin* promoter (blue arrow) followed by *eGFP* coding sequence (green box), and flanking *Ars* insulators (gray arrows). After co-injection of the donor vector with sgRNA for the donor vector, sgRNA for the genomic target site, and Cas9 mRNA, two patterns of insertion are predicted to occur due to NHEJ. (B) eGFP expression in *Gb-Ubx^KI-exon^*G_0_ stage 11 embryos. 4.7% of G_0_ embryos showed mosaic eGFP expression in the T3 legs (Table 3). (C) In G_1_ stage 14 embryos, the eGFP expression pattern was identical to that of the previously reported expression pattern of *Gb-Ubx* (Barnett et al., 2019; Matsuoka et al., 2015; Zhang et al., 2005). Asterisks in (B) and (C) mark position of embryonic head. (D) Wild type embryo does not show any eGFP expression. (E) G_2_ heterozygous mutants show an eGFP expression pattern identical to the pattern of *Gb-Ubx*. (F) G_2_ homozygous mutants showed strong eGFP expression, and also showed phenotypes characteristic of *Gb-Ubx^CRISPR^* mutants, including shortened T3 legs and formation of leg-like structures on the A1 segment (white arrowhead). (G) Assessment of knock-in event by using PCR and Sanger sequencing. We designed PCR primers specific for each putative junction (black arrows flanking areas **a** and **b** of panel (A)). All three homozygous G_2_ individual mutant animals assayed showed bands of the expected size for each junction. (H) Sequence analysis using the same primers indicated in panel A for genotyping confirmed that several deletions were generated due to NHEJ events at each junction. Black: genomic sequence; Blue: CRISPR target sequence; red: PAM sequence; pink: deleted nucleotides. Scale bar: 200 μm in (B)-(F). Embryonic staging as per (Donoughe and Extavour, 2016).

To confirm the integration of donor sequence into the genome, we performed PCR and sequence analysis. We designed specific primers for each 5’ and 3’ junction point (a and b in Fig. 3A). All three examined embryos showed the expected amplicon size for both junctions (Fig. 3G), suggesting that at least two copies of the donor vector fragment were integrated into the genome. Sequence analysis further confirmed the integration of the donor plasmid into the genome, and that indel mutations were generated at each junction (Fig. 3H). To determine how many copies of donor sequence were integrated into the genome, we performed copy number estimation by quantitative RT-PCR. The copy number of the donor plasmid was estimated as the copy ratio of the *eGFP* gene to that of the endogenous *orthodenticle* gene (*Gb-otd*), which is known to have only one copy in the genome (Nakamura et al., 2010; Ylla et al., 2021). The results of this analysis indicated that three copies of the donor plasmid were likely integrated into the genome (Supplementary Fig. 4A).

To determine whether the function of the target gene was indeed disrupted by this knock-in/knock-out strategy, we examined eGFP expression and morphology in G_2_ *Gb-Ubx^CRISPR-KI^* embryos. Among the G_2_ *Gb-Ubx^CRISPR-KI^* embryos, we found they displayed one of two different intensities of eGFP expression (Fig. 3E’ and 3F’). We determined their genotype by quantitative RT-PCR, and found that the individuals with strong eGFP expression were homozygous mutant, and the individuals with weak eGFP expression were heterozygotes (Supplementary Fig. 4B). Consistent with the genotyping result, crickets with weak eGFP expression displayed no detectable morphological abnormalities (Fig. 3E and 3E’). Crickets with strong eGFP expression, however, had smaller T3 legs and formed leg-like structures rather than pleuropodia on the A1 segment (Fig. 3F and 3F’) These phenotypes, which were the same as those observed in the *Gb-Ubx^CRISPR^* homozygous mutant (Fig. 2C and 2D), further suggested that the strong eGFP-expressing crickets were homozygous mutants.

### Knock-in of donor vector sequence at the Gb-abd-A locus

To confirm the efficiency and utility of this method, we next chose the Hox gene *Gb-abdominal-A* (*abd-A*) as a target (Table 1). We designed sgRNAs for the sequence within the exon just upstream of the homeodomain (Fig. 4A). We co-injected 50 ng/μl of sgRNA for the *Gb-abd-A* locus, 50 ng/μl of sgRNA for the donor vector, 100 ng/μl of Cas9 mRNA, and 100 ng/μl of donor vector into fertilized cricket eggs.

**Figure 4.**
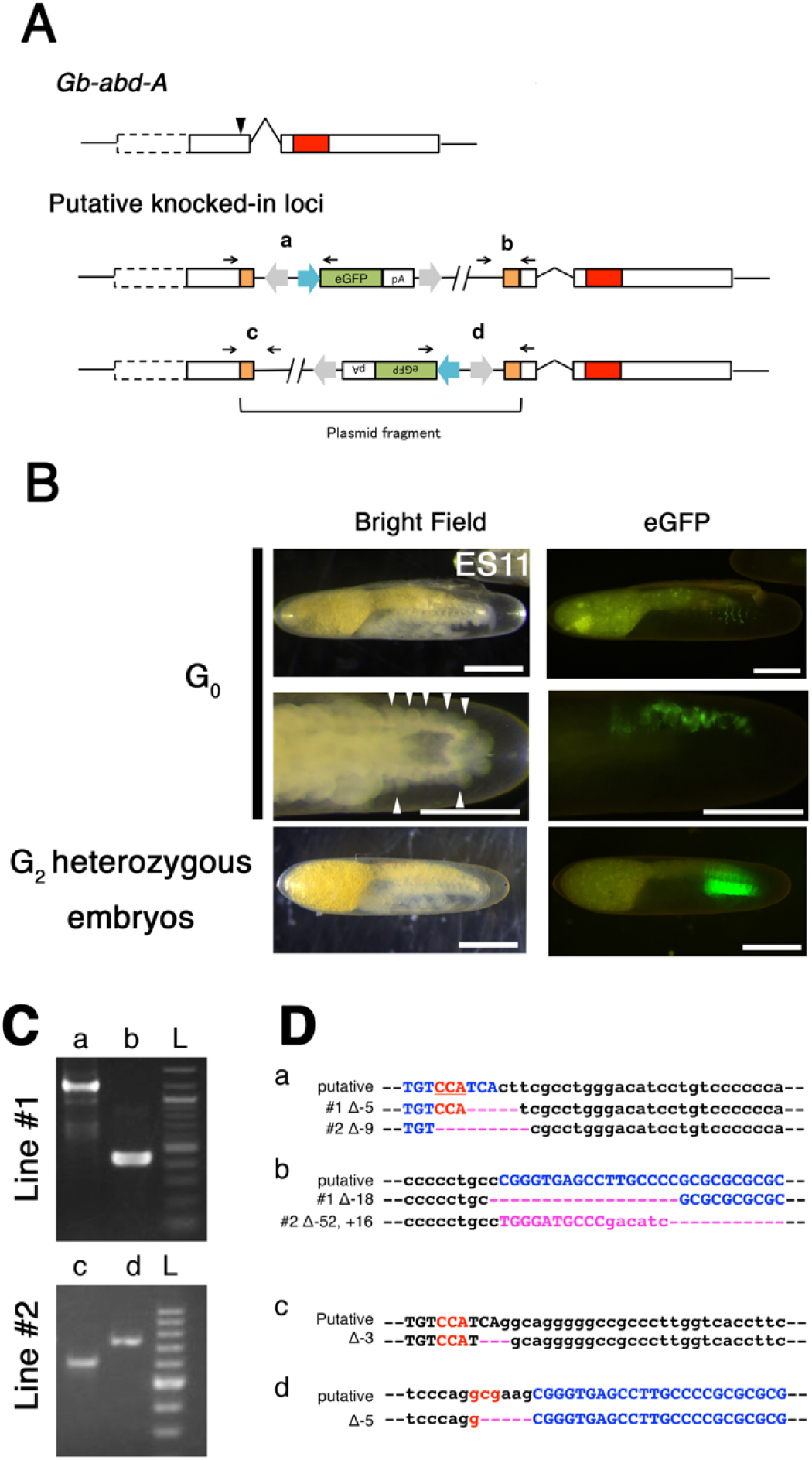
Exonic knock-in/Knock-out against *Gb-abd-A*. (A) Scheme of knock-in experiment against a *Gb-abd-A* exon. White box: exons; red box: homeodomain; black arrowhead: sgRNA target site. We used the same donor vector construct as that used in the experiment against *Gb-Ubx* (Fig. 3), substituting a *Gb-abd-A* exon-specific sgRNA. Two patterns of insertion are predicted to occur due to NHEJ. (B) Expression of eGFP in G_0_ and G_2_ *Gb-abd-A^KI-exon^*embryos. 10.6% of G_0_ *Gb-abd-A^KI-exon^* embryos showed mosaic eGFP expression in the abdomen of stage 11 embryos (Table 3). We found that eGFP expression was accompanied by ectopic phenotypic leg-like structures (white arrowheads), consistent with the loss of *Gb-abd-A* activity. In G_2_ *Gb-abd-A^KI-exon^* stage 11 embryos, the expression pattern of eGFP was identical to the previously reported expression pattern of *Gb-abd-A* (Barnett et al., 2019; Matsuoka et al., 2015; Zhang et al., 2005). (C) Assessment of knock-in events by using PCR and Sanger sequencing. Genomic DNA was extracted from homozygous G_2_ *Gb-abd-A^KI-exon^* stage 11 embryos and used as a PCR template. We designed PCR primers specific for each putative junction (black arrows and regions **a**, **b** and **c**, **d** in panel (A)). The expected amplicon size was detected for each junction. L = ladder. (D) Sequence analysis using the primers for genotyping confirmed that multiple deletions or insertions were generated, likely due to the NHEJ events at each junction. Black: genomic sequence; Blue: CRISPR target sequence; red: PAM sequence; pink: deleted or inserted nucleotides. Scale bar: 200 μm in (B). Embryonic staging as per (Donoughe and Extavour, 2016).

Of 38 injected G_0_ embryos, five showed mosaic eGFP expression in the abdomen (Fig. 4B). Four G_0_ adults were individually backcrossed with wild type counterparts of the opposite sex to obtain multiple G_1_ crickets. We obtained one stable transgenic line, in which eGFP expression in G_2_ embryos was similar to the previously documented expression pattern of *Gb-abd-A* transcript (Barnett et al., 2019; Matsuoka et al., 2015; Zhang et al., 2005) (compare Fig. 4B with Supplementary Fig. 1K). In a replicate injection experiment, we obtained a second such transgenic line (Table 3). PCR and sequence analysis confirmed that one of the two lines contained the plasmid fragment in the sense orientation, and the second line contained the plasmid fragment in the antisense orientation (Fig.4C). Copy number estimation analysis results suggested that a single plasmid fragment was integrated into the genome in each line (Supplementary Fig.4B).

In the *Gb-abd-A^KI-exon^* lines, eGFP expression was detectable in nymphs even through the cuticle (Supplementary Fig. 3C’, D’). We further detected eGFP expression in adult male and female internal organs (Supplementary Fig. 5). In wild type females, a pair of ovaries, each comprising hundreds of ovarioles, is located in the anterior abdomen (Nandchahal, 1972). Mature eggs are stored in an egg chamber at the posterior of each ovariole, and eggs are subsequently moved posteriorly through the oviduct. The posterior end of the oviduct is connected to the uterus, where fertilization takes place, located at the base of the ovipositor (Supplementary Fig. 5C’’). In *Gb-abd-A^KI-exon^*mutant females, eGFP expression was detected in the posterior portion of the oviduct (compare Supplementary Fig. 5B’ and B’’ to Supplementary Fig.5A’). We found, however, that the oviduct and uterus were not connected in *Gb-abd-A^KI-exon^*mutant females, and these females were not able to lay eggs. *Gb-abd-A^KI-exon^* mutant females also generated ectopic ovipositors (Supplementary Fig. 6U). In *Gb-abd-A^KI-exon^*males, ubiquitous eGFP expression was detected throughout the testis (compare Supplementary Fig. 5E’ and E’’ to Supplementary Fig. 5D’). The observed eGFP expression in females is reminiscent of the expression pattern of *D. melanogaster abd-A* in the developing female genital disc (which gives rise to the somatic reproductive structures including the oviduct in this fruit fly (Epper, 1983; Sánchez and Guerrero, 2001), and in the adult oviducts (Foronda et al., 2006).

We also detected eGFP expression at the anterior tip of the testis in *Gb-Ubx^KI-exon^* adult males (Supplementary Fig. 5G). *abd-A* expression has not, to our knowledge, been previously detected in the *D. melanogaster* male genital disc (Freeland and Kuhn, 1996), which gives rise to male somatic reproductive structures. However, high-throughput sequencing data from the modENCODE project do report *abd-A* expression in the adult *D. melanogaster* testis (Brown et al., 2014).

To confirm whether the eGFP expression observed in *Gb-abd-A^KI-exon^*and *Gb-Ubx^KI-exon^* animals reflects the endogenous expression of target genes in a given tissue, we performed quantitative RT-PCR on eGFP-positive and eGFP-negative tissues. In a previous study, we performed an RNA-seq analysis using *G. bimaculatus* adult brain and gonad tissue from both sexes (Whittle et al. 2021). In this dataset, we did not detect expression of *Gb-Ubx* nor *Gb-adb-A* in the brain (Fig. S7A, B). We therefore used brain as a negative control tissue for the expression of *Gb-Ubx* and *Gb-abd-A*. We found a clear correlation between *eGFP* and *Gb-abd-A* expression levels in the oviduct (eGFP-positive) and the uterus (eGFP-negative) in *Gb-abd-A^KI-exon^* females (Supplementary Fig. 7C and D). In the adult testis of *Gb-abd-A^KI-exon^* males, where we observed strong *eGFP* expression, we confirmed that the expression of *eGFP* and *Gb-abd-A* was also well correlated (Supplementary Fig. 7E). These results indicate that *eGFP* expression correlates well with *Gb-abd-A* expression, and are consistent with the interpretation that *eGFP* expression in these knock-in crickets reflects target gene expression.

In contrast, we observed no correlation between *Gb-Ubx* and *eGFP* expression levels in either eGFP-positive (anterior tip of testis) or eGFP-negative (rest of the testis) tissues, even though *eGFP* expression levels showed a clear correlation between those tissues (Supplementary Fig. 7F and G). It might be possible that *Gb-Ubx* is expressed at low levels throughout the ovary and that we were therefore unable to detect *eGFP* expression in this organ except for at the tips of ovaries. We found that the *Gb-Ubx* expression level in the wild type ovary and ovary tip was comparable to the level detected in the brain; in other words, it was effectively undetectable (Supplementary Fig. 7H). These results suggest that *Gb-Ubx* is not expressed, or is expressed only at levels undetectable in ou transcriptome (Whittle et al. 2021), at the tip of wild type ovaries, even though eGFP expression was detected there in *Gb-Ubx^KI-exon^* adults (Supplementary Fig. 5G). Taking into account the fact that three copies of the donor sequence were integrated in the *Gb-Ubx^KI-exon^* line (Supplementary Fig. 4), it might be possible that the observed eGFP expression in the tip of the ovary from *Gb-Ubx^KI-exon^*was an artifact due to the multiple copies of the expression cassette integrated at the *Gb-Ubx* locus, rather than being reflective of a true wild type *Gb-Ubx* expression domain. It is also possible that the *Gb-act* promoter used in the expression cassette might be more or differently active compared to the endogenous *Gb-Ubx* promoter. The fact that the *Gb-Ubx^KI-exon^*line did not show any abnormality in ovary development is consistent with this notion. Taken together, we conclude that it is possible to visualize target gene expression by knocking an expression cassette into a target gene locus, but careful verification should be done before making a conclusion as to all domains and levels of target gene expression.

### Targeted insertion of an expression cassette into an intron of the Gb-abd-A locus

Kimura and colleagues (Kimura et al., 2014) demonstrated that in *D. rerio* the homology independent method could be applied for trapping endogenous enhancer activity by inserting a donor sequence containing an expression cassette into the 5’UTR of genes of interest. We aimed to apply this technique to *G. bimaculatus* by attempting to knock-in a donor vector into the intronic region of *Gb-abd-A* (Fig. 5A).

**Figure 5.**
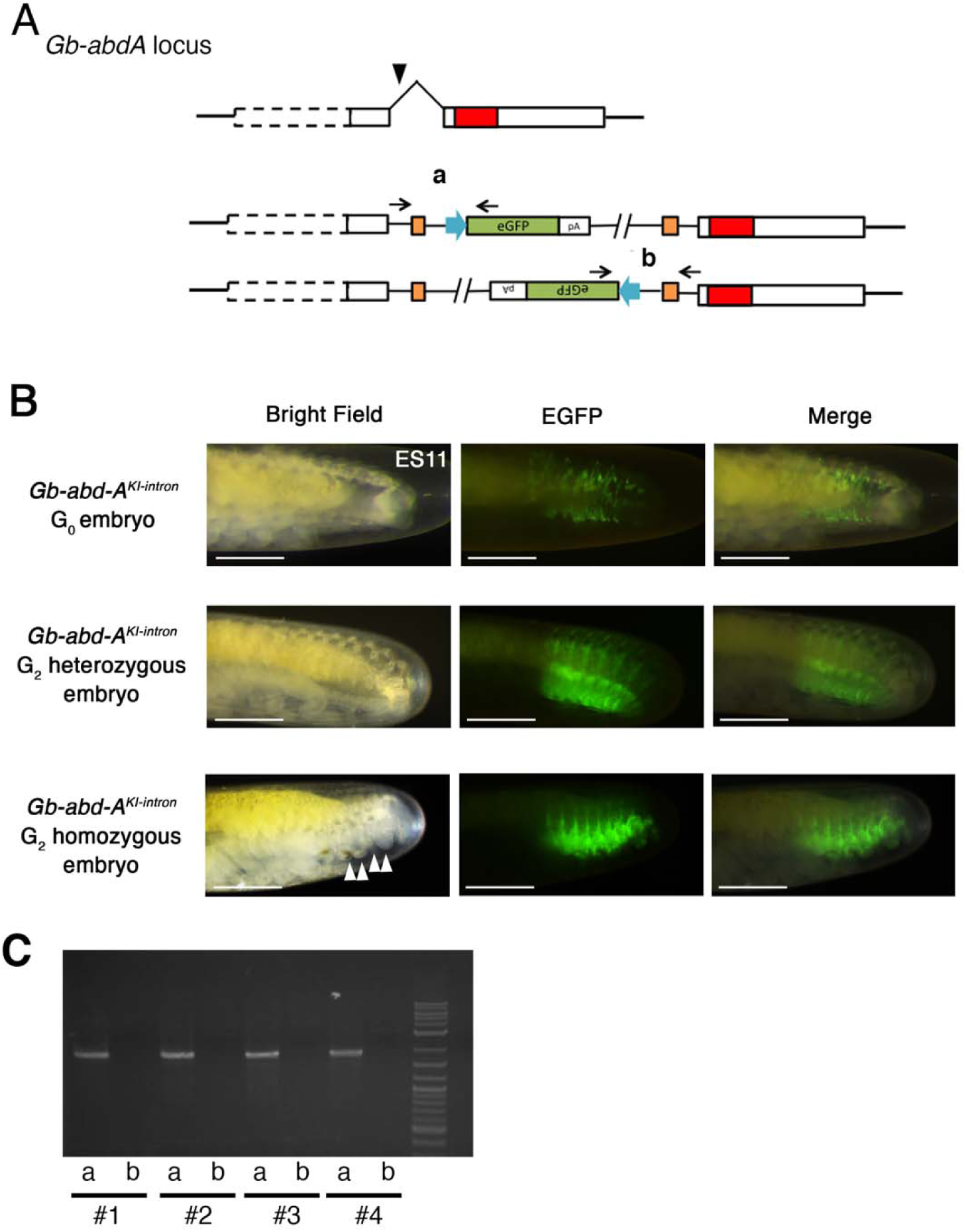
Intronic knock-in against *Gb-abd-A*. (A) Scheme of knock-in experiment targeted to a *Gb-abd-A* intron. White boxes: exons; red box: homeodomain; black arrowhead: sgRNA target site. We used the same donor vector construct as that used in the experiment against *Gb-Ubx* (Fig. 3), substituting a *Gb-abd-A* intron-specific sgRNA. Two patterns of insertion are predicted to occur due to NHEJ. (B) Expression pattern of eGFP in G_0_ and G_2_ stage 11 *Gb-abd-A^KI-intron^* embryonic abdomens. In *Gb-abd-A^KI-intron^*embryos, patchy eGFP expression was observed but embryos did not show the ectopic abdominal appendage phenotype that was observed in *Gb-abd-A^KI-exon^* embryos (Fig. 4B). *Gb-abd-A^KI-intron^* G_2_ heterozygous embryos show abdominal eGFP expression corresponding to the known pattern of embryonic *Gb-abd-A* transcripts (Barnett et al., 2019; Matsuoka et al., 2015; Zhang et al., 2005), and the embryos did not show any morphological abnormality. *Gb-abd-A^KI-intron^* G_2_ homozygous embryos generated ectopic leg-like structures on the abdomen (white arrowheads) as observed in G_0_ *Gb-abd-A^CRISPR^* embryos (compare with (B’)). (C) Assessment of knock-in event by using PCR with *Gb-abd-A^KI-intron^* G_2_ homozygous embryos. We designed PCR primers specific for each putative junction (black arrows and regions **a** and **b** in panel (A)). Scale bar: 500 μm in (B). Embryonic staging as per (Donoughe and Extavour, 2016).

We co-injected an sgRNA against an intron of *Gb-abd-A*, together with all other relevant reagents as described above (sgRNA against the donor vector, donor vector, and Cas9 mRNA) (Fig. 5A). Of 100 injected eggs, two eggs showed mosaic expression of eGFP in the abdomen (Fig. 5B). When the donor sequence was inserted into an exon in the previous experiment (see “*Knock-in of donor vector sequence at the Gb-abd-A locus*” above), eGFP expression was accompanied by a phenotype of ectopic leg-like structure development on abdominal segments (Fig.4B; Supplementary Fig. 6), as previously observed in *Gb-abd-A* RNAi experiments (Barnett et al., 2019). However, when the plasmid fragment was inserted into an intron, the region expressing eGFP did not generate ectopic leg-like structures in G_0_ embryos (Fig. 5B). This apparent absent or minimal loss-of-function phenotype in the intron knock-in embryos might explain the relatively high survival rate of the intron-targeted G_0_ embryos (28% of injected G_0_ embryos survived to adulthood) compared to that of the exon-targeted G_0_ embryos (4.9% to 8.5% of 41 and 47 injected embryos survived to adulthood; Table 3).

We obtained one *Gb-abd-A^KI-intron^* line, in which we confirmed that one donor vector sequence was likely integrated into the target region in a forward orientation (Fig. 5C). We carefully inspected the morphology of the *Gb-abd-A^KI-intron^* adult crickets to assess whether potential post-embryonic functions of the target gene were affected by insertion of the donor sequence into an intron. G_1_ heterozygous *Gb-abd-A^KI-intron^*females did not show the supernumerary ovipositors observed in G_1_ heterozygous *Gb-abd-A^KI-exon^* female adults (Supplementary Fig .6W compared with 6U), and they laid eggs normally (data not shown). For further confirmation, we examined the morphology of G_2_ homozygous *Gb-abd-A^KI-intron^* mutants. Approximately 25% of examined G_2_ eggs showed strong eGFP expression, which we interpret as likely indicative of a homozygous mutant. All G_2_ embryos with strong eGFP expression generated leg-like structures on the abdomen (Fig. 5B), suggesting that the function of the target gene was somewhat affected in the homozygous condition, unlike in the mosaic condition exhibited by G_0_ embryos (Fig. 5B). Thus, even though the knock-in into an intron somewhat affected target gene expression. G_0_ embryos with intronic insertions did not show abnormalities, unlike the embryos with exonic insertions. This increases the developmental success of intronic knock-in embryos, and accordingly, the possibility of establishing a transgenic line.

## Discussion

In the present study, we demonstrated that targeted knockout and knock-in by using the CRISPR/Cas9 system works efficiently in the cricket *G. bimaculatus*. We performed functional analysis of CRISPR/Cas9-induced mutations in the Hox genes *Gb-Ubx* and *Gb-abd-A* during embryogenesis and at post-embryonic stages. We found that the cleavage efficiency of the CRISPR/Cas9 system was much higher than that previously reported for artificial nucleases in this cricket (Watanabe et al., 2012; Watanabe et al., 2014). We demonstrated that gene knock-in via a homology-independent method is effective in this cricket, and successfully applied it to functional analysis of Hox genes by knocking a donor sequence into an exon of the target gene to disrupt the function of the target gene (knock-in/knock-out). In addition, we succeeded in trapping endogenous gene activity using this method and revealed a number of new expression domains that had not been previously observed with traditional methods (Barnett et al., 2019; Matsuoka et al., 2015; Zhang et al., 2005). This homology-independent method is technically simpler than the homology-dependent methods, as the donor plasmid does not need to be newly made for each target region.

### CRISPR/Cas9 system vs RNA interference

Delivering the proper amount of genome editing constructs at the proper time is important for efficient outcomes. For example, gene editing in embryos injected around 3h AEL, at which stage energids are distributing throughout the yolk, was highly efficient, but both detectable mosaic phenotypes (of *Gb-lac2* crispants) and the rate of NHEJ decreased with later injections at blastoderm stages (Supplementary Fig. 2B, D, and H). Thus, we suggest that optimization of delivery conditions can be achieved by using pigmentation genes as an index.

In our study, reproducibility and severity of phenotypes generated with CRISPR/Cas9 were greater than those obtained with RNAi (Fig. 2D and 2F). We speculate that the efficiency of RNAi-mediated knockdown may be influenced by when and at what levels the target gene is expressed. In the case of *Gb-abdA* and *Gb-Ubx*, these genes are expressed much later in development (embryonic stage 5, about 1.5 days after egg laying) than the stage at which we performed injections (within 3 hours after egg laying) (Matsuoka et al., 2015, Donoughe et al., 2016. Correspondingly, much higher concentrations of dsRNA proved necessary to produce even mild phenotypes (5-6 μg/μl; Fig. 2E), than those typically used for RNAi against most genes in this cricket (1-2 ug/ul; e.g. Donoughe et al., 2014). In contrast, indel mutations generated at earlier stages by genome editing techniques resulted in clearly detectable, severe phenotypes (Supplementary Fig. 2). Several studies have demonstrated that genome editing techniques can sometimes be adequate for functional analysis of target genes in mosaic G_0_ individuals (e.g. Daimon et al., 2015; Martin et al., 2016; Matsuoka and Monteiro, 2018). However, it is often difficult or impossible to unambiguously identify mutant cells in such mosaics. In this regard, establishment and maintenance of stable mutant lines as performed herein, allows for less ambiguous phenotypic analysis in this cricket.

Although the CRISPR/Cas9 system is efficient, it offers little to no conditionality, which can complicate study of the many genes that act pleiotropically during development (Minelli, 2016). For example, in the case of *Gb-abd-A*, the gene acts to repress leg formation in the abdomen at embryonic stages (Supplementary Fig. 6), while at adult stages, it regulates proper development of female genitalia (Supplementary Fig. 7). Likely because of this latter phenotype, we were unable to obtain homozygous *Gb-abdA^CRISPR^*animals. To overcome this problem, sophisticated genetic methods, like balancer chromosomes (Miller et al., 2019), will need to be developed in the future. In this regard, RNAi offers more options for conditional control of gene function. By controlling the timing of injection of dsRNA, target gene activity can be knocked down at any desired developmental stage in *G. bimaculatus* (Dabour et al., 2011; Nakamura et al., 2008; Takahashi et al., 2009). Thus, while the CRISPR/Cas9 system is a powerful new tool for gene function analyses, RNAi remains a useful technique for this system.

### Application of homology-independent knock-in method for functional analysis of endogenous genes

Homology-independent knock-in methods will expand our ability to analyze the function of target genes in this hemimetabolous insect model. Here, we demonstrated one such application, the KI/KO method, which allows for isolation of mutants without PCR-based genotyping. When analyzing mutant phenotypes, affected individuals must typically be distinguished either by their morphology or by molecular methods to detect changes in target gene product levels or functions. In the case of the *Gb-Ubx* mutant, we would have needed to distinguish subtle differences in the T3 and A1 embryonic segments (Fig. 2C), requiring destructive sampling. Moreover, antibodies against target genes may not be routinely available in many cases. The KI/KO method allows us to distinguish mutant individuals based on marker gene expression. Even heterozygous and homozygous mutants can sometimes be distinguished based on the intensity of marker gene expression. A similar strategy was employed in mosquitos via HDR (McMeniman et al., 2014). In the present study, we could easily identify the eGFP expression resulting from the KI/KO event because it matched the previously characterized expression pattern for *Gb-Ubx* (Barnett et al., 2019; Matsuoka et al., 2015; Zhang et al., 2005). However, for target genes with previously uncharacterized expression domains, analysis may be more complex.

The promoter used in all expression cassettes herein is the same one used in a previous study to drive ubiquitous constitutive expression (Nakamura et al., 2010). Nevertheless, all knocked-in lines showed an eGFP expression pattern that was spatially and temporally restricted like that of the target gene. We speculate that the promoter in the expression cassette acts as a minimal promoter, and that the observed eGFP expression results from trapping endogenous enhancer activity. The eGFP expression was not caused by the fusion to the endogenous gene product, since both the line containing an inverted orientation of the donor sequence, and the knock-in line targeting an intronic region, showed similar eGFP expression patterns. To enhance the usefulness of this method, identification and use of a ubiquitous and strong promoter could in principle drive exogenous marker gene expression in the whole embryo without being subject to positional effects.

A remarkable feature of the homology-independent knock-in method is the length of sequence that can be integrated. In case of knock-in through HDR, a few kb of sequence can be integrated into the genome in arthropods (Gilles et al., 2015; McMeniman et al., 2014). In this study, through NHEJ, at least six kb of plasmid sequence was integrated into the genome. Furthermore, in some cases, three copies of plasmid sequence were integrated in tandem into the genome (Supplementary Fig. 4). In this case, we speculate that first the donor plasmids were digested and combined via NHEJ, and then the combined fragment was knocked in into the genome via NHEJ, suggesting that homology-independent knock-in might be able to integrate several tens of kb of sequence into the genome. For example, one study showed that a 200 kb BAC vector could be integrated into a rodent genome through a similar strategy (Yoshimi et al., 2016). This method might therefore be used for direct functional comparison of genomic regions by exchanging homologous regions between related species of interest.

The efficiency of knock-in through NHEJ is high, but to improve its feasibility as a technique for functional genetic analysis in this cricket, future studies may be able to further enhance efficiency by optimizing at least one of two parameters, as follows: (1) *Enhance expression level of a fluorescent cassette*. Empirically, G_0_ crickets showing mosaic eGFP expression tend to transmit their knocked-in transgene to their offspring. To increase the efficiency of obtaining knock-in lines, future efforts should therefore focus on increasing the number of mosaic marker gene expression cassettes in G_0_ embryos. We also observed that some expression cassettes seemed to show higher expression levels than others and were therefore easier to detect in mosaic G_0_s. Inclusion of inducible expression elements such as a heat shock promoter or a modified Gal4/UAS system might help to enhance the activity of the expression cassette. However, enhancement of fluorescent cassette expression may cause artificial or ectopic expression as we observed in the ovaries of *Gb-Ubx^KI-exon^* (Supplementary Fig. 5G’’). (2) *Introduce insulator sequences into the donor cassette*. Positional effects might also in principle prevent the full potential activity of the expression cassette. In this study, our vector plasmid contained insulators of the sea urchin *Hemicentrotus pulcherrimus* arylsulfatase gene (Takagi et al., 2011) on either side of the expression cassette (see Materials and Methods), but we nonetheless detected eGFP expression in a pattern matching that of the target gene. To achieve more effective insulation, future studies might evaluate several different combinations of insulator orientations, which can affect insulator activity (Tchurikov et al., 2009). Alternatively, other insulators such as that of the *gypsy* retrotransposon (Modolell et al., 1983) might be additional options for future optimization (Carballar-Lejarazú et al., 2013).

In conclusion, we provide evidence that the CRISPR-Cas9 system works well for both knock-in and knock-out in the cricket *G. bimaculatus*. In depth analysis of CRISPR-Cas9-induced mutations revealed optimized injection timing. In addition, we succeeded in the targeted functional knock-in of exogenous sequences into the genome through NHEJ, resulting in expression-tag reporter lines for target genes.

## Materials and Methods

### Cricket husbandry

All adult and juvenile *Gryllus bimaculatus* were reared in plastic cages at 26–30 °C and 50% humidity under a 10-h light, 14-h dark photoperiod. They were fed on artificial fish food (Tetra) or Purina cat food (item #178046). For microinjections, fertilized eggs were collected on a wet kitchen towel in a plastic dish and incubated at 28 °C as previously described (Barry et al., 2019; Watanabe et al., 2017).

### Construction of sgRNA vectors

sgRNA target sequences were designed and their off-target sites were predicted with the CasOT program (Xiao et al., 2014). From the suggested candidates, we selected target sequences having high GC content (around 70%), beginning with guanidine for efficient *in vitro* transcription by the T7 promoter, and with off-target sites containing at least 3 mismatches as per Ren and colleagues (Ren et al., 2014). Two synthetic oligonucleotides (5’-ATAG-N_19_-3’ and 5’-AAA-N_20_) were annealed and inserted into the BsaL site of the modified pDR274 vector (Addgene plasmid #42250) to expand its utility (the GGN_18_NGG sequence was present in the original pDR274, while in the modified vector, a GN_19_NGG sequence was used). We confirmed insertion by Sanger sequence analysis.

### Synthesis of sgRNA and mRNA

For sgRNA synthesis, the template for *in vitro* transcription was digested from the vectors generated as described above with DraⅠ. For Cas9 mRNA synthesis, the template for *in vitro* transcription was digested from pMLM3613 (Addgene catalogue #42251) with PmeⅠ. Both sgRNA and Cas9 mRNA were *in vitro* transcribed using mMESSAGE mMACHINE T7 Kit (Life Technologies catalogue #AM1344), and purified by ethanol precipitation. For the Cas9 mRNA, we attached a poly-A tail by using a poly-A tailing Kit (Life Technologies catalogue #AM1350). The concentration of synthesized RNAs was estimated by NanoDrop and gel electrophoresis.

### Construction of donor plasmids

The *eGFP*bait-2A-RFP donor plasmid was generated in a pUC57 vector by commercial artificial composition (GeneScript). The *DsRed*bait-*G’act*-*eGFP* donor plasmid was generated based on the *eGFP*bait-2A-RFP donor plasmid. First, 2A-RFP was digested using BglⅡ and NotⅠ. *Gb-act*-*eGFP* was also digested from a pXL-BacⅡ-G’act-*eGFP* vector and ligated to generate the *eGFP*bait-G’act-*eGFP* vector. Then, we digested this *eGFP*bait vector using BglⅡ and SacⅠ. We amplified *DsRed*bait with primers (5’ to 3’) DsRed_fwd: GCTCAGATCTCTTGGAGCCGTACTGGAAC, and DsRed_rev: GTACGAGCTCCATCACCGAGTTCATGCG. The amplicon was ligated to generate the *DsRed*bait-G’act-*eGFP* donor plasmid. The *DsRed*bait-2×Ars_rev-G’act-*eGFP*-2×Ars_fwd donor plasmid was generated based on the *DsRed*bait-G’act-*eGFP* donor plasmid. The Ars insulator sequence ArsInsC from *H. pulcherrimus* (Takagi et al., 2011) was amplified from an ArsInsC-containing plasmid (kind gift of Assoc. Prof. Naoaki Sakamoto, Hiroshima University, Japan) and integrated on either side of the expression cassette in the donor plasmid.

### Microinjection

Cas9 mRNA, sgRNA, and donor vectors were injected into 2-5h AEL cricket eggs. Cricket eggs were aligned in a groove 0.7 mm deep and 0.7 mm wide made with 2% agarose in 1x phosphate-buffered saline (PBS) using a custom mold as previously described (Barry et al., 2019; Watanabe et al., 2017) and filled with 1xPBS. Needles for injection were made by pulling glass capillaries with filament (Narishige catalogue # GD-1) with a pipette puller (Sutter Instrument catalogue # P-1000IVF), using the following pulling program: (1) x3 Heat; 858, Pull; 0, Velocity; 15, Time; 250, Pressure; 500, and (2) x1 Heat; 858, Pull; 80, Velocity; 15, Time; 200. To minimize the invasiveness of the injection, the tips of the pulled needles were sharpened and ground to a 20° angle by using a Micro Grinder (Narishige catalogue # EG-400). Approximately 5 nl of solution was injected into eggs with a Micro Injector (Narishige catalog # IM300). After injection, eggs were moved to a fresh Petri dish and submerged in fresh 1xPBS containing 50 U/ml penicillin and 50 μg/ml streptomycin (15070-063, Thermo Fisher), and incubated at 28°C. During the incubation period, the 1xPBS with penicillin and streptomycin was replaced every day. We observed fluorescent protein expression at the stages when the target gene was known to be expressed. Genomic DNA was extracted from 7 d AEL eggs and adult T3 legs and used for insertion mapping and sequence analyses. After two days of incubation, injected cricket eggs were moved to wet filter paper in a fresh Petri dish for hatching.

### Detection of indel mutations

After Cas9 nuclease digests a target sequence, the disrupted sequence is repaired by either the NHEJ or the HDR cell machinery. To confirm a KO mutation, we searched for errors repaired by the NHEJ pathway, which sometimes introduces or deletes nucleotides at the digested site during the repair process. Since disruption or repair are unlikely to take place identically in all cells of an injected G_0_ embryo, G_0_ animals are expected to contain heterogeneous sequences at the CRISPR targeted site, and thus to be heterozygous for a putative Cas9-induced indel. To confirm the activity of the sgRNAs, the Surveyor^TM^ Mutation Detection Kit (Transgenomic,) was used. This assay relies on a “surveyor” nuclease that can recognize and digest a heteroduplex DNA structure. First, genomic DNA was extracted from whole eggs or part of the T3 leg by a phenol chloroform method as previously described (Barry et al., 2019; Watanabe et al., 2017). Subsequently, approximately 200 bp of the targeted region was amplified by PCR from genomic DNA (Table 4). PCR conditions were optimized to reduce non-specific amplification or smearing. To create the putative heterogeneous DNA structure for the nuclease assay, the PCR product was heated to 98°C for five minutes, and then re-annealed by gradually cooling down to 30°C. Half of the PCR product was digested with the Surveyor^TM^ nuclease, and the other half was used as a negative control and incubated without the nuclease. Digestion was confirmed by agarose gel electrophoresis. For sgRNAs that yielded indels in the target sequence, digest of the PCR product by the Surveyor^TM^ nuclease is expected to produce split fragments around the CRISPR targeted site relative to the negative control; the latter should not be digested by the nuclease and thus should remain intact and run at the same size as the original amplicon. Positive PCR products were extracted from the gel, purified with the QIAquick Gel Extraction Kit (Qiagen catalogue #28506), and sub-cloned into the pGEM-Teasy vector (Promega catalogue #A1360) using TA-cloning. The vectors were used for Sanger sequence analysis.

**Table 4.**
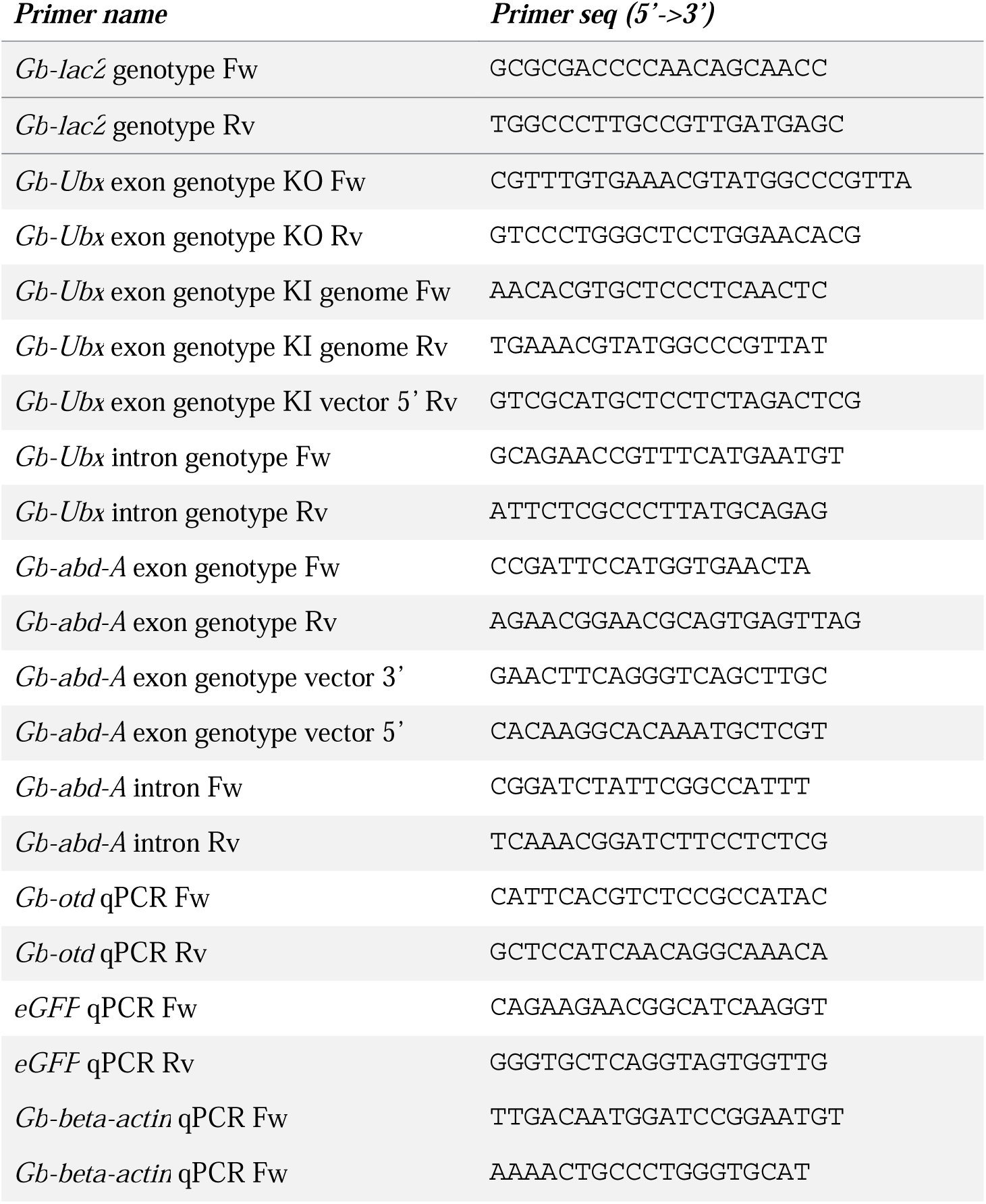
Primers used in this study.

### Amplicon sequence analysis

After a 1h egg collection, eggs were incubated for the desired length of time at 28°C. We co-injected 0.5 μg/μl sgRNA and 1 μg/μl Cas9 mRNA into fertilized cricket eggs after each of these incubation periods. Five days after injection, genomic DNA was extracted individually from three eggs from each of the four tested injection times; the latter analysis was performed in biological triplicate. Amplicon sequence analysis was performed by using MiSeq (Illumina), and the preparation of DNA libraries and sequencing reactions were performed according to the manufacturer’s instructions. We read ∼10,000 reads for on-target regions and ∼50,000 reads for off-target regions. We chose off-target sites that were most similar to the target site relative to other sequences in the genome. As we mentioned above about the criteria for selecting target site, the off-target site has three mismatch sequences compared to the target site. The assembly of output paired end reads was performed by using CLC Genomic Workbench (CLC Bio, QIAGEN Digital Insights). The relative proportions of reads containing indels and substitutions in the individual eggs were calculated with the online-tool CRISPResso (Pinello et al., 2016). We used the Integrative Genomic Viewer (Broad Institute) for investigation of the distribution of indels and substitutions (Thorvaldsdóttir et al., 2013).

### Insertion mapping

Genomic DNA was extracted from eGFP-positive eggs of each line. Due to the specifics of this knock-in method, two types of insertion of vector fragment (sense and antisense orientations) would be expected to occur; we therefore performed PCR using primers designed against either side of the putative junction. PCR was performed using target region-specific (upstream or downstream of sgRNA recognition site) and donor vector-specific primers (sequence within *eGFP* for forward integration and M13Fw for reverse integration). Primer sequences are listed in Table 4. Positive PCR products were extracted from the gel, purified by using the QIAquick Gel Extraction Kit (Qiagen catalogue #28506), and sub-cloned into the pGEM-Teasy vector (Promega catalogue #A1360) using TA-cloning. The vectors were used for Sanger sequence analysis.

### Embryo fixation, whole mount in situ hybridization, and immunohistochemistry

Embryos were dissected in 1xPBS and fixed with 4% paraformaldehyde in 1xPBS + 0.1% Tween (PBT) for 1h at 4℃. The fixed embryos were dehydrated stepwise in 25%, 50%, 75%, and 100% methanol in 1xPBT with five minutes per wash. The dehydrated embryos were stored in 100% methanol at -30℃. Whole-mount *in situ* hybridization with digoxigenin (DIG)-labeled antisense RNA probes was performed as previously described (Niwa et al., 2000; Zhang et al., 2005). Immunohistochemistry was performed as follows: Fixed embryos were rehydrated stepwise in 75%, 50%, and 25% solutions of methanol/ PBT and finally in 100% PBT for five minutes in each solution. After blocking with 1% bovine serum albumin (BSA) (Thermo Fisher) in PBT for one hour at room temperature, embryos were incubated with an anti-UbdA antibody FP6.87 (Kelsh et al., 1994) (Developmental Studies Hybridoma Bank) diluted 1:200 in 1% BSA/PBT overnight at 4℃. After washing with PBT three times, embryos were incubated in 1% BSA/PBT for one hour at room temperature, and then incubated with Alexa Fluor 488-conjugated Goat Anti-mouse IgG(H+L) (Invitrogen catalogue #A32723) diluted 1:400 in 1% BSA/PBT for one hour at 4℃. After washing the embryos with PBT once for 10-60 minutes, embryos were counter-stained with DAPI (Sigma catalogue #10236276001) stock solution 1mg/mL diluted 1:1000 in PBT for ten minutes, and then washed with PBT two times for 10-60 minutes per wash. PBT was then substituted with 25% and 50% glycerol/PBT to clear embryos for microscopy.

### Copy number estimation and detection of endogenous Hox gene expression in eGFP-positive tissues by using quantitative RT-PCR

To estimate the number of plasmid fragments integrated into the genome via NHEJ events, we performed quantitative RT-PCR using genomic DNA from individual 5 day-old embryos of wild type, *Gb-Ubx^KI-exon^*, and *Gb-abd-A^KI-exon^* lines, and compared relative quantity values of the inserted *eGFP* gene to an endogenous gene *Gb-otd*, which is present in a single copy in the *G. bimaculatus* genome (Ylla et al., 2021). Genomic DNA was extracted from embryos using Cica geneus® Total DNA Prep kit (for Tissue) (Kanto Chemical), according to the manufacture’s protocol. Real-time quantitative PCR was performed using the power SYBR Green PCR Master Kit (Applied Biosystems catalogue #4368577) and an ABI 7900 Real Time PCR System (Applied Biosystems) as described previously (Nakamura et al., 2008). Primer sequences are listed in Table 4.

To examine whether the eGFP expression observed in gonads of *Gb-Ubx^KI-exon^*and *Gb-abd-A^KI-exon^* reflects the endogenous expression of *Gb-Ubx* or *Gb-abd-A* in those tissues, we performed quantitative RT-PCR using total cDNA from tissues with or without eGFP expression. We dissected out gonads and brain in 1xPBS, and then transferred the dissected tissues to RNA later (Thermo Fisher). Total mRNA was extracted with RNeasy MinElute Cleanup Kit (Qiagen), according to the manufacture’s protocol. Equal amounts of total RNA were used for reverse transcription reaction with SuperScript III Reverse Transcriptase (ThermoFisher). Quantitative RT-PCR was performed with THUNDERBIRD® SYBR® qPCR Mix (TOYOBO) and LightCycler 96 (Roche). The expression levels of *eGFP, Gb-Ubx* and *Gb-abd-A* were normalized to the level of *Gb-beta-tubulin.* Primer sequences used were as previously described (Barnett et al., 2019).

## Acknowledgements

We thank N. Sakamoto for the Ars plasmid, and the members of the Extavour and Mito labs for discussion.

## Competing Interests

No competing interests declared.

## Funding

This work was supported by National Science Foundation award number IOS-1257217 and funds from the Howard Hughes Medical Institute and Harvard University to CGE, an Overseas Research Fellowship (grant # 693) from the Japan Society for the Promotion of Science (JSPS) to TN, a Grant-in-Aid for Young Scientists B (grant # JP16K21199, JP26870415) from the JSPS to TW, and a Grant-in-Aid for Scientific Research (B) (grant # JP26292176) from the JSPS to TM.

## Data availability

Not applicable.

## Notes

### Competing Interest Statement

The authors have declared no competing interest.

### Summary of Updates

we have added new supporting data to the supplement, and clarifying methodological and interpretative text to the main manuscript body.

