## Supplementary Material for "Establishment of CRISPR/Cas9-based knock-in in a hemimetabolous insect: targeted gene tagging in the cricket *Gryllus bimaculatus*"

The Supplementary Information for this study consists of the following:

- Supplementary Results and Discussion
- Supplementary Figure Legends S1 through S7
- Supplementary References
- Supplementary Figures S1 through S7

#### Supplementary Results and Discussion

*Analysis of Gb-Ubx*$^{CRISPR}$ *embryo phenotype*

To understand the genetic basis of the T3 phenotypes in the *Gb-Ubx*$^{CRISPR}$ homozygous mutants, we examined the embryonic expression pattern of the leg patterning genes *wingless* (*Gb-wg*), *decapentaplegic* (*Gb-dpp*), and *dachshund* (*Gb-dac*). Wild type T3 legs express *Gb-Ubx* (Barnett et al., 2019; Matsuoka et al., 2015; Zhang et al., 2005) and have T3 leg-specific expression of some leg-patterning genes. Namely, *Gb-wg* is expressed on the ventral side of each leg segment (Niwa et al., 2000) and on the dorsal side of the femur in the T3 leg (Supplementary Fig. 1A''), but not in the T1 or T2 legs (Supplementary Fig. 1A-A'). In *Gb-Ubx*$^{CRISPR}$ mutants, dorsal T3 femoral expression of *Gb-wg* was undetectable (Supplementary Fig. 1B''), consistent with a transformation of this appendage towards an anterior fate. *Gb-dpp* is expressed in several dorsal and ventral spots in wild type developing limbs, and in the T3 leg, *Gb-dpp* is expressed in five circumferential bands (Supplementary Fig. 1C; Niwa et al., 2000). In contrast, in *Gb-Ubx*$^{CRISPR}$ embryos, *Gb-dpp* expression in the T3 leg resembled the expression pattern observed in wild type T1 or T2 legs (Supplementary Fig. 1D'' and D'''' compared with Supplementary Fig. 1C). Taken together, the T3 leg-specific pattern of multiple leg-patterning genes was absent, consistent with disruption of the *Gb-Ubx* locus. These results suggest that the T3 segment in *Gb-Ubx*$^{CRISPR}$ mutants acquired characteristics of the T2 segment, consistent with a homeotic transformation loss-of-function phenotype of *Gb-Ubx*$^{CRISPR}$.

The appendage on the A1 segment, called the pleuropodium, is a transient appendage structure only observed during mid-embryogenesis (Rathke, 1844; Wheeler, 1892), and is thought to be involved in secreting hatching enzymes (Konopová et al., 2020; Slifer, 1937). In wild type embryos, *Gb-Ubx* is strongly expressed in this organ at early embryonic stages (Barnett et al., 2019; Matsuoka et al., 2015; Zhang et al., 2005), as are *Gb-Dll* and *tramtrack* (*Gb-ttk*) (Fig. 2E and Supplementary Fig. 1E and E'; Barnett et al., 2019). In *Gb-Ubx*$^{CRISPR}$ mutants, the pleuropodia appeared as small, twisted leg-like structures (Fig. 2E, Supplementary Fig. 1B''', D''', and H''') that lacked *Gb-ttk* expression (Supplementary Fig. 1F and F') and ectopically expressed *Gb-wg*, *Gb-dpp, Gb-Dll*, and *Gb-Dac* in patterns like those observed in wild type thoracic legs (Fig. 2E, Supplementary Fig. 1B''', D''', and H''').

We speculate that knock-out of *Gb-Ubx* may cause misexpression of other Hox genes and contribute to the *Gb-Ubx*$^{CRISPR}$ phenotype. To test this hypothesis, we examined the expression patterns of *Antennapedia* (*Antp*) and *abdominal-A* (*abd-A*) in *Gb-Ubx*$^{CRISPR}$ embryos. In wild type embryos *Gb-Antp* is not expressed in the pleuropodia (Supplementary Fig. 1I and I'), while *Gb-abd-A* is expressed in the posterior half of the A1 to A9 segments but not in the pleuropodia (Supplementary Fig. 1K and K'). In *Gb-Ubx*$^{CRISPR}$ embryos, the expression pattern of *Gb-abd-A* was unchanged (Supplementary Fig. 1L and L'), but *Gb-Antp* was misexpressed in the pleuropodia (Supplementary Fig. 1J and J'). We interpret these expression patterns as evidence that *Gb-Ubx* represses *Gb-Antp* expression in the pleuropodia, and the interruption of the *Gb-Ubx* locus causes pleuropodia to be transformed towards thoracic leg identity, which may be induced by misexpression of *Gb-Antp.*

*Analysis of Gb-abd-A mutants*

We detected several different phenotypes in *Gb-abd-A*$^{KI\text{-}exon}$ $G_0$ and $G_1$ mutants (Supplementary Fig. 5 and 6): (1) supernumerary leg-like structures on the abdomen; (2) fusion of cuticle segments on the abdomen; (3) additional ovipositors in heterozygous females. Supernumerary leg-like structures were observed on the abdomen of mosaic $G_0$ embryos and nymphs. To elucidate the genetic mechanism underlying the production of these structures, we examined the expression patterns of leg patterning genes in homozygous *Gb-abd-A*$^{KI\text{-}exon}$ embryos.

We found that *Gb-wg* was ectopically expressed along the ventral side of the supernumerary leg-like structures, as in wild type developing limbs, generated on the A8 and A9 (Supplementary Fig. 6C and D), and the pattern was retained at later stages (Supplementary Fig. 6G and H). *Gb-Dll* was ectopically expressed in the supernumerary leg-like structures in the A2-A9 segments (Supplementary Fig. 6K and L), which is similar to the expression pattern observed in the thoracic legs (Supplementary Fig. 6I). As the embryo developed, *Gb-Dll* expression was no longer detected in the supernumerary leg-like structures in the A2 throughA7 segments, but it retained detectable in the supernumerary leg-like structures in the A8 and A9 segments (Supplementary Fig. 6O and P). Taken together, misexpression of *Gb-Dll* led to the generation of leg-like structures on the early embryonic abdomen, while at later stages, the only structures expressing *Gb-wg* retained the expression of *Gb-Dll*. The strongly affected embryos did not hatch, suggesting that the low hatching rate observed in these $G_0$ embryos (Table 3) might be due to this phenotype. Similar appendage-like structures were also observed on the abdomen of the nymphs (Supplementary Fig. 6T).

The CRISPR/Cas9 system allowed us to efficiently produce severe mutants for *Gb-Ubx* and *Gb-abd-A*, which gave us the opportunity to investigate leg patterning in *G. bimaculatus*. The genetic mechanism for leg specification and patterning has been well studied in *D. melanogaster*, and two different mechanisms for regulating *Dll* expression are important for determining the segments that possess ventral appendages. One of the mechanisms is the activation of *Dll* gene by the segment polarity genes *wingless* (*wg*) and *engrailed* (*en*) (Cohen, 1990). Although *wg* and *en* are expressed in all segments, *Dll* expression is excluded from the abdominal segments in *D. melanogaster*. The repression of *Dll* in the abdomen is achieved by the Hox genes *Ubx*, *abd-A*, and *Abd-B* (Vachon et al., 1992). Lewis and colleagues (2000) addressed the relationship between *en* and Hox genes in the beetle*, Tribolium castaneum*. In that study, *T. castaneum abd-A* mutants showed ectopic pleuropodia throughout the abdomen resulting from ectopic *Dll* expression, and *en* expression was expanded to the distal region in the ectopic pleuropodia, indicating that *abd-A* represses both *Dll* and *en* expression in the abdomen of this beetle. In our cricket study, *Gb-abd-A*$^{KI\text{-}exon}$ mutants produced ectopic ventral appendages in the abdomen, and the ectopic appendages on the A8 and A9 are more like a leg-like structure (Supplementary Fig. 6D, H, L and P), which was not observed in *T. castaneum abd-A* mutants (Lewis et al., 2000). *Gb-Dll* was expressed in the ectopic ventral appendages throughout the abdomen, but ectopic *Gb-wg* expression was only observed in the leg-like structures in A8 and A9. We speculate that in A8-A9, disruption of *Gb-abd-A* leads to the misexpression *Gb-Dll* and results in the production of ectopic ventral appendages throughout the abdomen. In A8 and A9, *Gb-wg* was also derepressed due to the loss of *Gb-abd-A* activity, and probably concomitant expression of *Gb-wg* and *Gb-Dll* produced a leg like structure. It might be possible that *Gb-Abd-B* expression in A8 and A9 is not able to repress *Gb-wg* in this cricket. Similar misexpression of *Gb-wg* was observed in the ectopic A1 legs of *Gb-Ubx*$^{CRISPR}$ embryos. These results suggest that both *Gb-Ubx* and *Gb-abd-A* repress *Gb-wg* and *Gb-Dll* expression in the abdomen, that *Gb-Dll* expression alone is not sufficient, and that *Gb-wg* expression is necessary, for thoracic leg development.

*Potential roles of Gb-abd-A in insect genitalia development*

Supernumerary ovipositors were observed in $G_0$ and $G_2$ heterozygous *Gb-abd-A*$^{KI\text{-}exon}$ females (Supplementary Fig. 6U). To evaluate potential effects on internal abdominal organs, we dissected adult males and females. The ovipositor and the subgenital plate are located on the A8 segment in wild type females (Supplementary Fig. 6R). In *Gb-abd-A*$^{KI\text{-}exon}$ heterozygous females, the subgenital plate on the ventral side of the A9 was absent; instead, additional ovipositors were detected (compare Supplementary Fig. 6U to Supplementary Fig. 6R). Some *Gb-abd-A*$^{KI\text{-}exon}$ heterozygous females did not lay eggs probably due to this phenotype. We dissected *Gb-abd-A*$^{KI\text{-}exon}$ heterozygous females to assess potential effects on internal organs and found that many eggs were stored inside the egg chamber (Supplementary Fig. 5B), suggesting that egg production was not affected. We found that the posterior tip of the oviducts had not fused with the uterus (Supplementary Fig.6B' and B"). In *D. melanogaster iab-4* mutants, one of the *cis*-regulatory regions of *abd-A* affects gonadal development in females (Cumberledge et al., 1992). In severe cases, these mutants failed to form a gonad, and in mild cases, the gonad was formed but the ovary and oviduct junctions were not attached, and thus egg transfer from ovary to the uterus was blocked (Cumberledge et al., 1992). Taken together, these results suggest that the function of *Gb-abd-A* in the development of the adult ovary, specifically in ensuring appropriate joining of the uterus to the oviducts to allow egg release, is conserved between *D. melanogaster* and *G. bimaculatus*.

The genitalia have been considered as serially homologous to the ventral appendages, since several leg-patterning genes contribute to genitalia development in *D. melanogaster* (Estrada and Sanchez-Herrero, 2001). *D. melanogaster Abd-B* mutants transform male and female genitalia towards leg fate, and the transformation is accompanied by the ectopic expression of *Dll* and *Dac*, which are normally required in the legs (Estrada and Sanchez-Herrero, 2001). *D. melanogaster abd-A* also contributes to the development of the genitalia in female, since some *D. melanogaster abd-A* mutants lack or show abnormality in the ovaries (Foronda et al., 2006). However, the additional ovipositors generated in cricket *Gb-abd-A*$^{CRISPR}$ mutants are reminiscent of the phenotype of *D. melanogaster Abd-B* loss of function mutants, although the ectopically generated appendage in the cricket was an ovipositor rather than a leg. We speculate that *Gb-abd-A* represses some genes related to the outgrowth of appendages, potentially including *Gb-Dll*, during development of the inner genitalia. Under this hypothesis, when the activity of *Gb-abd-A* is reduced or disrupted, the generated outgrowth might be defined as an ovipositor by the action of *Gb-Abd-B*.

#### Supplementary Figure Legends

**Supplementary Figure S1. Expression pattern of limb patterning genes in *Gb-Ubx*$^{CRISPR}$ homozygous mutants.**

Expression pattern of limb patterning genes in wild type and *Gb-Ubx*$^{CRISPR}$ homozygous mutants. (A-A''') In wild type embryos, *Gb-wg* was expressed along the ventral side of appendages, and also in the dorsal femur in the T3 limb bud (arrowhead in A'') but not in the T1 or T2 limb buds. (B-B''') In *Gb-Ubx*$^{CRISPR}$ homozygous mutants, the dorsal expression domain of *Gb-wg* in the T3 limb bud was undetectable (B''). In pleuropodia, *Gb-wg* expression was only observed at the proximal region (B'''). (C) In wild type limb buds, *Gb-dpp* is expressed in each limb segment, and also in a broad expression domain in the femur and tibia of T3 but not T1 or T2 limb buds (arrowheads). (D-D'''') In *Gb-Ubx*$^{CRISPR}$ homozygous mutants, the T3-specific femur and tibia expression domains of *Gb-dpp* were undetectable (D'' and D''''). In pleuropodia, *Gb-dpp* was expressed as in the distal tip region of wild type limb buds (D'''). (E and E') *Gb-ttk* was expressed in the developing pleuropodia in wild type embryos. (F and F') In *Gb-Ubx* homozygous mutants, *Gb-ttk* expression was undetectable in the pleuropodia. (G-G''') In wild type embryos, *Gb-Dac* was expressed in the prospective femur and tibia of limb buds, but not in the pleuropodia. (H-H''') In *Gb-Ubx*$^{CRISPR}$ homozygous mutants, the expression pattern in the thoracic limb buds was not affected, but the pleuropodia showed *Gb-Dac* gene expression like that observed in limb buds (H'''). (I-I') In wild type embryos *Gb-Antp* was expressed in the thoracic limb buds, but not in the pleuropodia. (J-J') In *Gb-Ubx*$^{CRISPR}$ homozygous mutants, *Gb-Antp* was ectopically expressed in the pleuropodia. (K-K') In wild type embryos Gb-abd-A was expressed in the abdomen. (L-L') The *Gb-abd-A* expression pattern was not affected in *Gb-Ubx*$^{CRISPR}$ homozygous mutants. Scale bar: 200 µm. Embryonic staging as per (Donoughe and Extavour, 2016).

**Figure S2. In-depth analysis of the feature of mutagenesis by CRISPR/Cas9 system in the crickets.**

(A) Schematic of injection timeline. After a 1h egg collection and 1h subsequent incubation, eggs were further incubated for 1h, 3h, 5h and 9h, respectively before injection. (B, C) NHEJ mutation rate (indels) at both on-target and off-target sites at the *Gb-Ubx* locus. The NHEJ mutation rate decreased with increasing age of injection and was <1.3% at the studied off-target site. (D, E) NHEJ mutation rate (indels) at both on-target and off-target sites at the *Gb-abd-A* locus NHEJ mutation rate decreased with increasing age of injection and was <1.2% at the studied off-target site. (F, G) Pattern of NHEJ mutations at each injection time point in the sequence around the CRISPR target site. Black bar indicates the site targeted for CRISPR/Cas9-induced double-stranded breaks. The pattern of NHEJ mutations was similar for all injection ages, but the rate decreased with increasing age of injection. (H-H''') Cuticular phenotype of mosaic $G_0$ hatchlings resulting from injection with *Gb-Lac2* sgRNA (Fig. 1) at each tested injection time point.

**Supplementary Figure S3. eGFP expression in the KI crickets.**

(A, A') eGFP expression in $G_1$ *Gb-Ubx*$^{KI\text{-}exon}$ heterozygous male nymphs was visible in the hindwings, which are on the T3 segment. (B, B') *Gb-Ubx*$^{KI\text{-}exon}$ heterozygous adults also showed eGFP expression in T3 legs. (C,C') In $G_0$ mosaic *Gb-abd-A*$^{KI\text{-}exon}$ nymphs, eGFP expression was detected in patchy cells of the abdominal cuticle. (D, D') In $G_1$ *Gb-abd-A*$^{KI\text{-}exon}$ nymphs, eGFP expression was observed in the cuticle of A1 to A8, which is similar to the previously reported embryonic expression pattern of *Gb-abd-A* (Barnett et al., 2019; Matsuoka et al., 2015; Zhang et al., 2005). (E,E') Higher magnification of region boxed in white in (D, D').

**Supplementary Figure S4. Expression levels and genomic copy number of the integrated eGFP genes in the KI crickets.**

(A) Relative quantity of *eGFP* mRNA in embryos of *Gb-abd-A*$^{KI\text{–}exon}$ and *Gb-Ubx*$^{KI\text{-}exon}$ lines. Embryos of each KI line were divided into two sample groups, respectively, according to the intensity of eGFP fluorescence (eGFP weak/strong). *Gb-beta-actin* was used as an internal standard. n = 5 for each sample group. (B) Estimation of genomic copy numbers of the transgene *eGFP* in the KI individuals by quantitative RT-PCR. The single copy gene *Gb-otd* was used as the reference. The results may indicate that the *Gb-abd-A*$^{KI\text{–}exon}$ line has one copy of the integrated plasmid, and that the *Gb-Ubx*$^{KI\text{–}exon}$ line has three copies of the integrated plasmid. eGFP weak- and strong-expressing embryos in (A) corresponded to heterozygous and homozygous genotypes for the *eGFP* insertion, respectively.

**Supplementary Figure S5. eGFP expression in genitalia of KI crickets.**

(A-C) Adult female viscera in ventral view made visible by removing the cuticle. (A, A') Ventral posterior view of WT adult female abdomen, indicating cerci (ci) and single ovipositor (ov). (B-B") Dissection revealed that eGFP expression was observed at the posterior tips of the oviducts, and the oviducts were not fused with the uterus (B"). (C, C') In *Gb-abd-A*$^{KI\text{-}intron}$ females, eGFP expression in the oviducts was brighter and broader than that detected in *Gb-abd-A*$^{KI\text{-}exon}$ females (B'), but the oviducts were fused with the uterus (C"). (D-F) Adult male viscera in ventral view made visible by removing the cuticle. (E-F) Single testes dissected out of the abdominal cavity. In *Gb-abd-A*$^{KI\text{-}exon}$ and *Gb-abd-A*$^{KI\text{-}intron}$ $G_2$ adult males, ubiquitous eGFP expression was observed in the testis (E' and F'). (G-G") *Gb-Ubx*$^{KI\text{-}exon}$ females showed eGFP expression at the anterior tip of the ovaries. (H and H') In *Gb-Ubx*$^{KI\text{-}exon}$ males, no eGFP expression was detected in the testis. Scale bar: 2 mm.

**Supplementary Figure S6. Developmental and morphological phenotypes in *Gb-abd-A*$^{KI\text{-}exon}$ mutants.**

(A, B) Expression pattern of *Gb-wg* in ES9 wild type embryos. (C, D) Expression pattern of *Gb-wg* in ES9 homozygous *Gb-abd-A*$^{KI\text{-}exon}$ $G_2$ embryos. An ectopic thoracic leg-like pattern of *Gb-wg* was observed in the ectopic leg-like structures generated on A8 and A9. (E, F) Expression pattern of *Gb-wg* in ES13 wild type embryos. (G, H) Expression pattern of *Gb-wg* in ES13 homozygous *Gb-abd-A*$^{KI\text{-}exon}$ $G_2$ embryos. *Gb-wg* expression was still detected in the ectopic leg-like structure on A8. (I, J) Expression pattern of *Gb-Dll* in ES9 wild type embryos. (K, L) Expression pattern of *Gb-Dll* in ES9 homozygous *Gb-abd-A*$^{KI\text{-}exon}$ $G_2$ embryos. *Gb-Dll* was expressed in all ectopic leg-like structures generated on the abdomen, but only in A8 did *Gb-Dll* show an expression pattern similar to the pattern in wild type developing thoracic limb buds. (M, N) Expression pattern of *Gb-Dll* in ES13 wild type $G_2$ embryos. (O, P) Expression pattern of *Gb-Dll* in ES13 homozygous *Gb-abd-A*$^{KI\text{-}exon}$ $G_2$ embryos. *Gb-Dll* expression was only detected in the leg-like structure on the A8. (Q, R) Wild type nymph abdomen. (S) Approximately 10% of $G_0$ *Gb-abd-A*$^{KI\text{-}exon}$ nymphs showed fusion of abdominal cuticle in some segments (white brackets). (T) Approximately 10% of $G_0$ *Gb-abd-A*$^{KI\text{-}exon}$ nymphs showed leg-like structures on the abdomen. (U) $G_1$ heterozygous *Gb-abd-A*$^{KI\text{-}exon}$ female nymphs showed ectopic ovipositors (arrowheads). (V) eGFP expression in $G_1$ heterozygous *Gb-abd-A*$^{KI\text{-}exon}$ nymphs. (W) *Gb-abd-A*$^{KI\text{-}}$

$^{intron}$ nymphs never developed ectopic ovipositors. (X) eGFP expression in $G_1$ *Gb-abd-A*$^{KI\text{-}intron}$ nymphs. Scale bars: 500 µm in (A) through (P); 2 mm in (Q) through (T). Embryonic staging as per (Donoughe and Extavour, 2016).

**Supplementary Figure S7. Relative quantification of *eGFP* expression level in the KI lines.**

**Supplementary Figure S8. Knock-in against *Gb-Ubx* intronic region.**
(A) Scheme of knock-in experiment targeted to a *Gb-Ubx* intron. White boxes: exons; red box: homeodomain; black arrowhead: sgRNA target site. We used the same donor vector construct as that used in the experiment against *Gb-Ubx* (Fig. 3), substituting a *Gb-Ubx* intron-specific sgRNA. Two patterns of insertion are predicted to occur due to NHEJ. (B, B') Expression pattern of eGFP in $G_2$ *Gb-Ubx*$^{KI\text{-}exon}$ stage 17 embryos. (C, C') Expression pattern of eGFP in $G_2$ *Gb-Ubx*$^{KI\text{-}intron}$ stage 17 embryos. *Gb-Ubx*$^{KI\text{-}exon}$ embryos (B') showed shorter legs than $G_2$ *Gb-Ubx*$^{KI\text{-}intron}$ embryos (C'). Scale bars: 500 µm. Embryonic staging as per (Donoughe and Extavour, 2016).

#### Figure S1

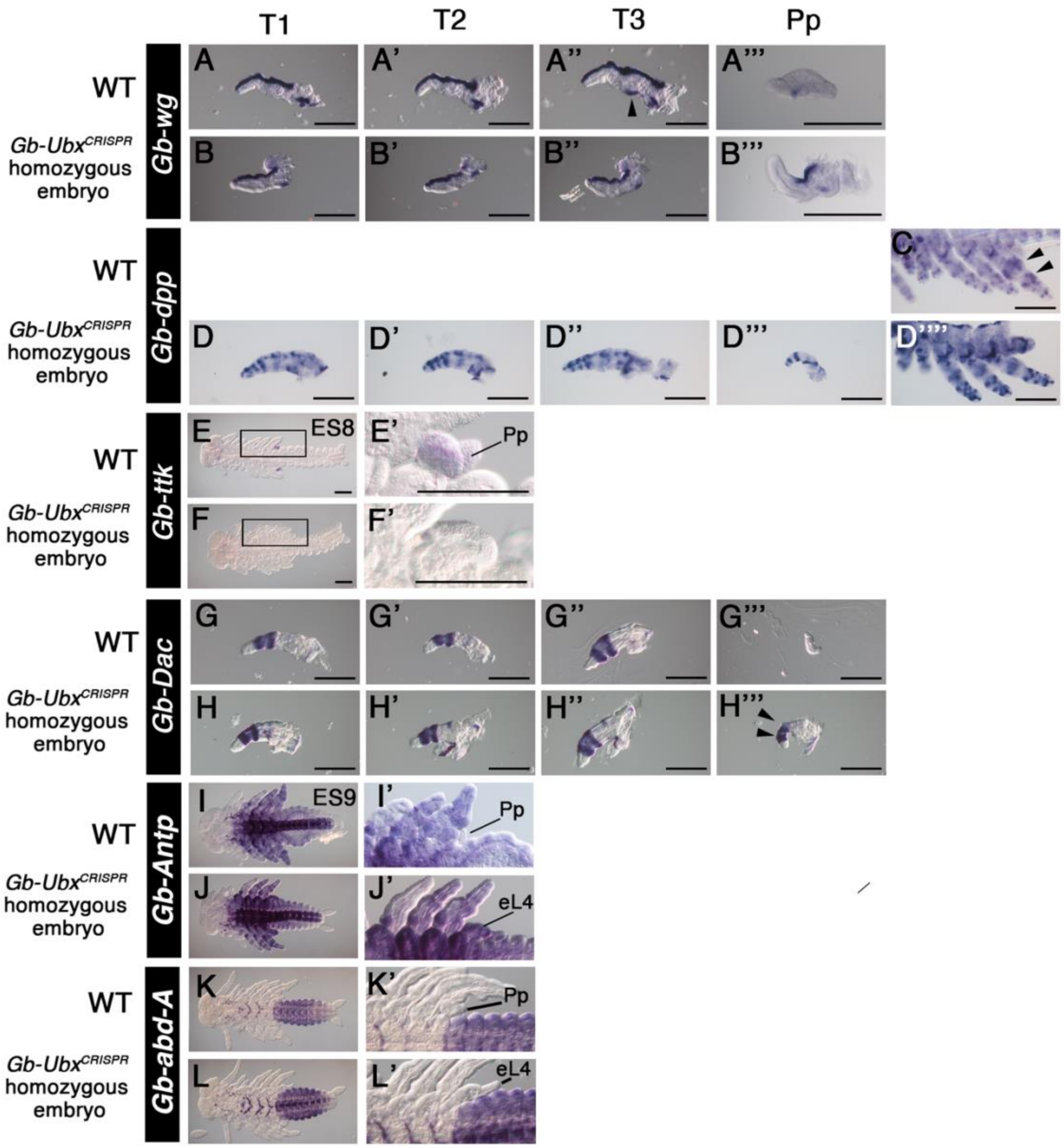

T1
T2
T3
Pp
WT
Gb-Ubx^CRISPR homozygous embryo
Gb-wg
Gb-dpp
Gb-ttk
Gb-Dac
Gb-Antp
Gb-abd-A
A
A'
A''
A'''
B
B'
B''
B'''
C
D
D'
D''
D'''
D''''
E
ES8
E'
Pp
F
F'
G
G'
G''
G'''
H
H'
H''
H'''
I
ES9
I'
Pp
J
J'
eL4
K
K'
Pp
L
L'
eL4

**Figure S2**

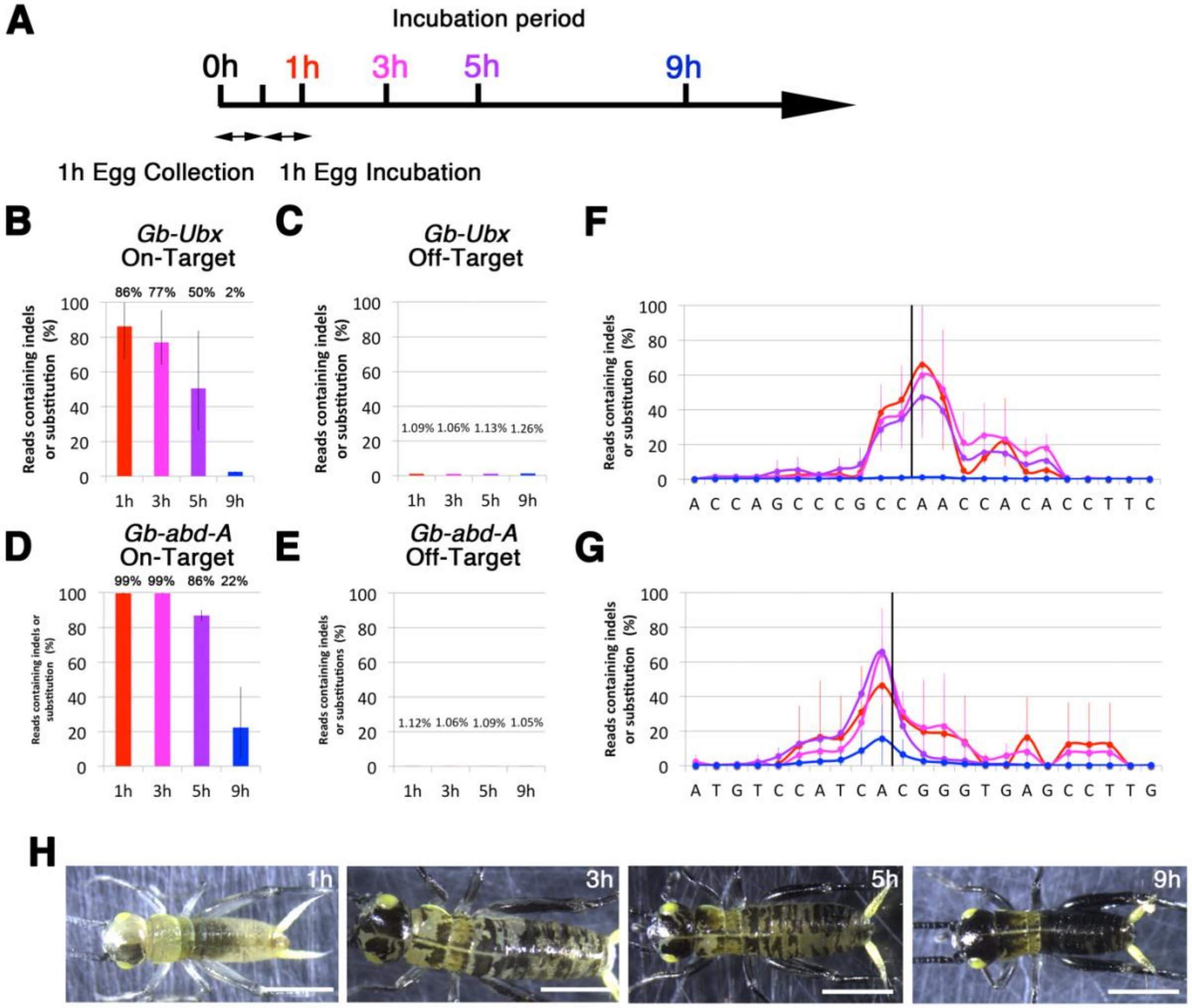
A
Incubation period
0h
1h
3h
5h
9h
1h Egg Collection
1h Egg Incubation
B
Gb-Ubx
On-Target
86% 77% 50% 2%
Reads containing indels or substitution (%)
1h 3h 5h 9h
C
Gb-Ubx
Off-Target
1.09% 1.06% 1.13% 1.26%
F
A C C A G C C C G C C A A C C A C A C C T T C
D
Gb-abd-A
On-Target
99% 99% 86% 22%
E
Gb-abd-A
Off-Target
Reads containing indels or substitutions (%)
1.12% 1.06% 1.09% 1.05%
G
A T G T C C A T C A C G G G T G A G C C T T G
H
1h
3h
5h
9h

**Figure S3**

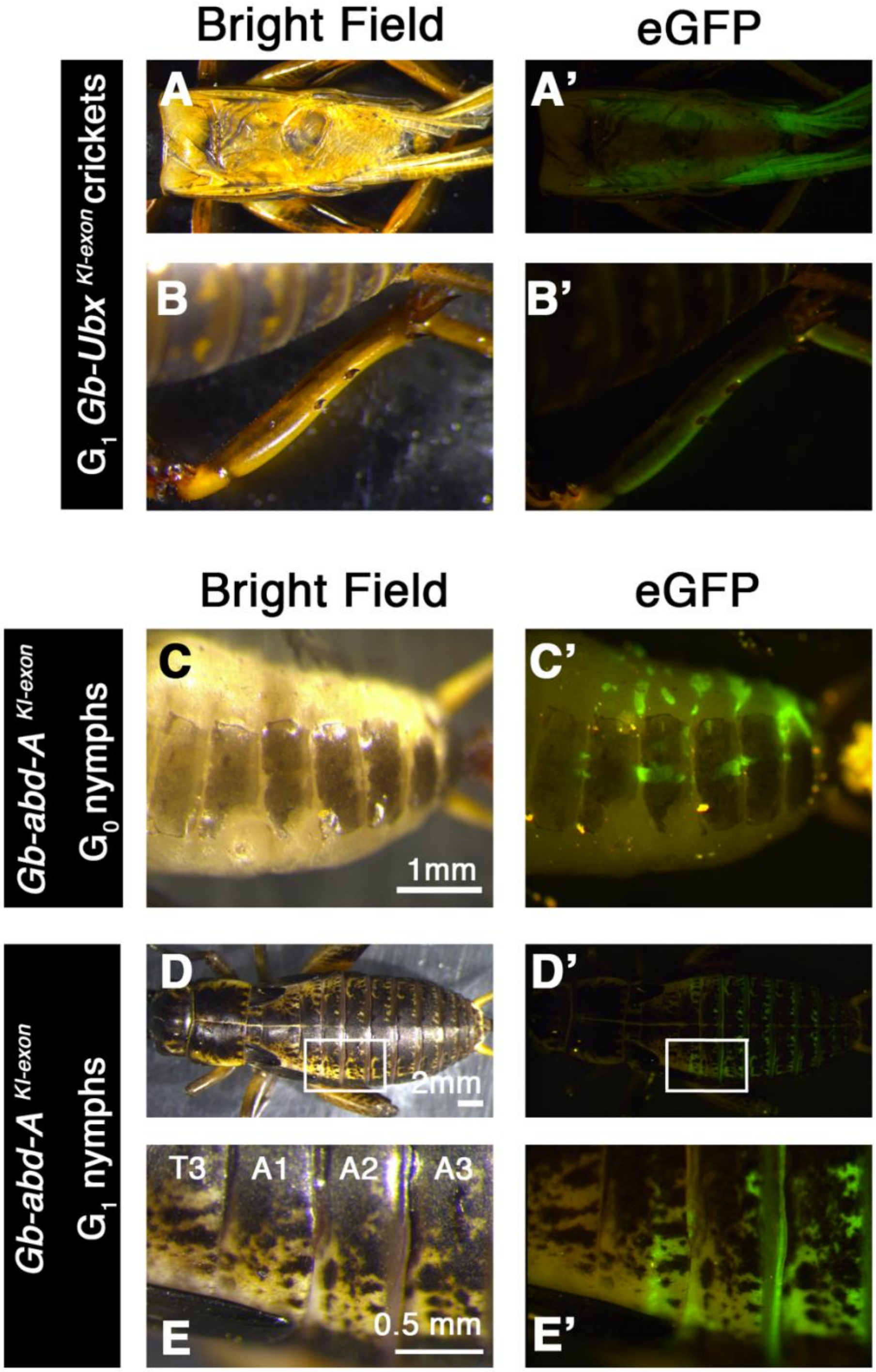
Bright Field
eGFP
$G_1$ *Gb-Ubx* $^{KI-exon}$ crickets
A
A'
B
B'
Bright Field
eGFP
*Gb-abd-A* $^{KI-exon}$ $G_0$ nymphs
C
C'
1mm
*Gb-abd-A* $^{KI-exon}$ $G_1$ nymphs
D
D'
2mm
T3
A1
A2
A3
E
E'
0.5 mm

**Figure S4**

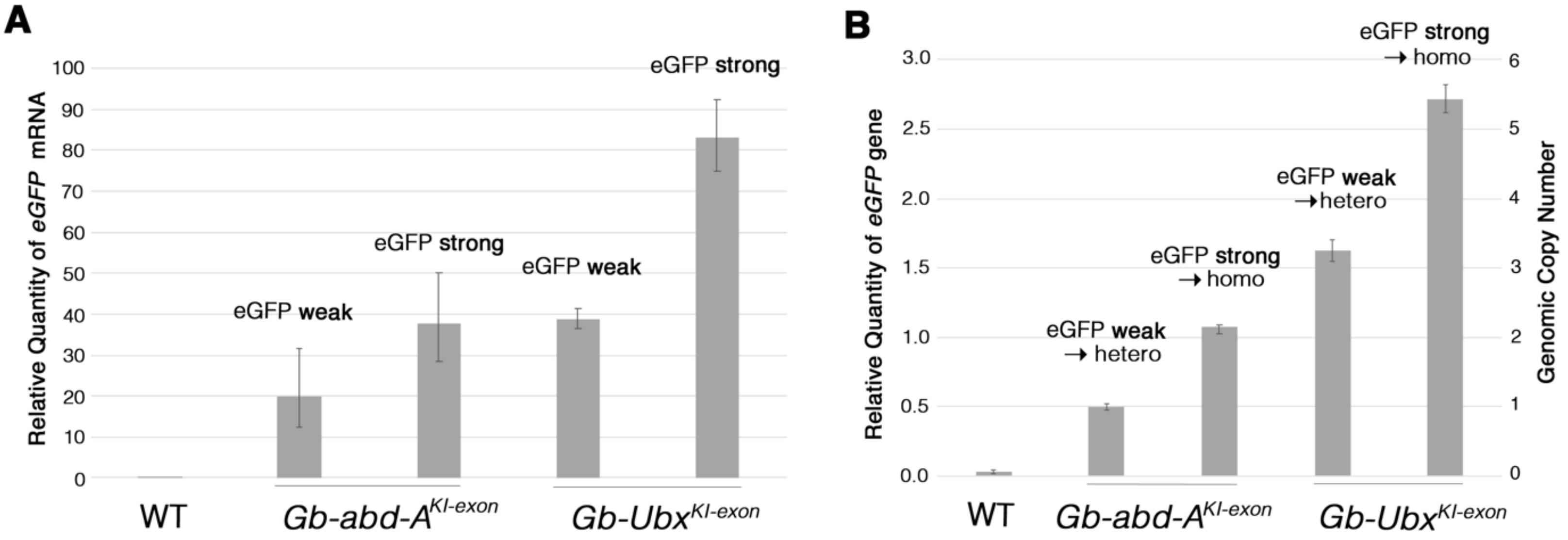
A
Relative Quantity of eGFP mRNA
100
90
80
70
60
50
40
30
20
10
0
eGFP weak
eGFP strong
eGFP weak
eGFP strong
WT
Gb-abd-A^KI-exon
Gb-Ubx^KI-exon
B
Relative Quantity of eGFP gene
3.0
2.5
2.0
1.5
1.0
0.5
0.0
Genomic Copy Number
6
5
4
3
2
1
0
eGFP weak
→ hetero
eGFP strong
→ homo
eGFP weak
→hetero
eGFP strong
→ homo
WT
Gb-abd-A^KI-exon
Gb-Ubx^KI-exon

#### Figure S5

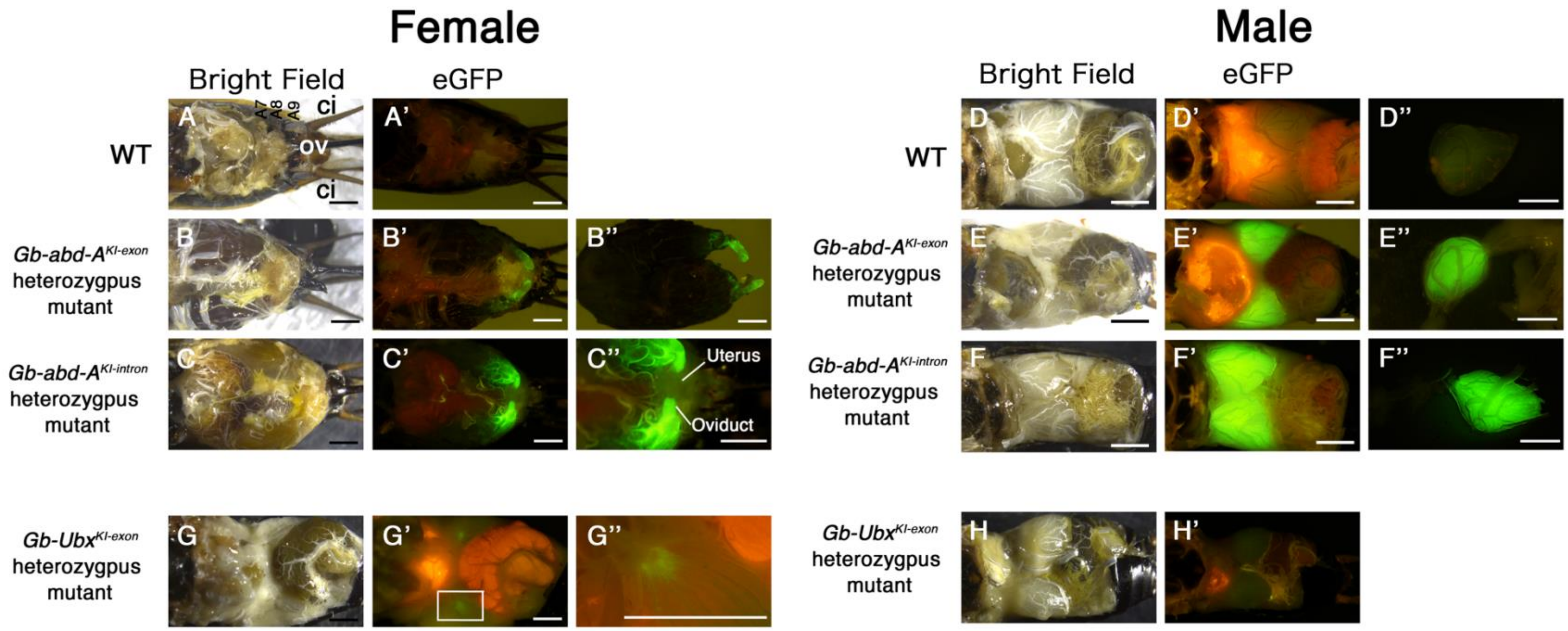

Female
Male
Bright Field
eGFP
Bright Field
eGFP
WT
A
A7
A8
A9
ci
ov
ci
A'
Gb-abd-A^KI-exon heterozygpus mutant
B
B'
B''
Gb-abd-A^KI-intron heterozygpus mutant
C
C'
C''
Uterus
Oviduct
WT
D
D'
D''
Gb-abd-A^KI-exon heterozygpus mutant
E
E'
E''
Gb-abd-A^KI-intron heterozygpus mutant
F
F'
F''
Gb-Ubx^KI-exon heterozygpus mutant
G
G'
G''
Gb-Ubx^KI-exon heterozygpus mutant
H
H'

#### Figure S6

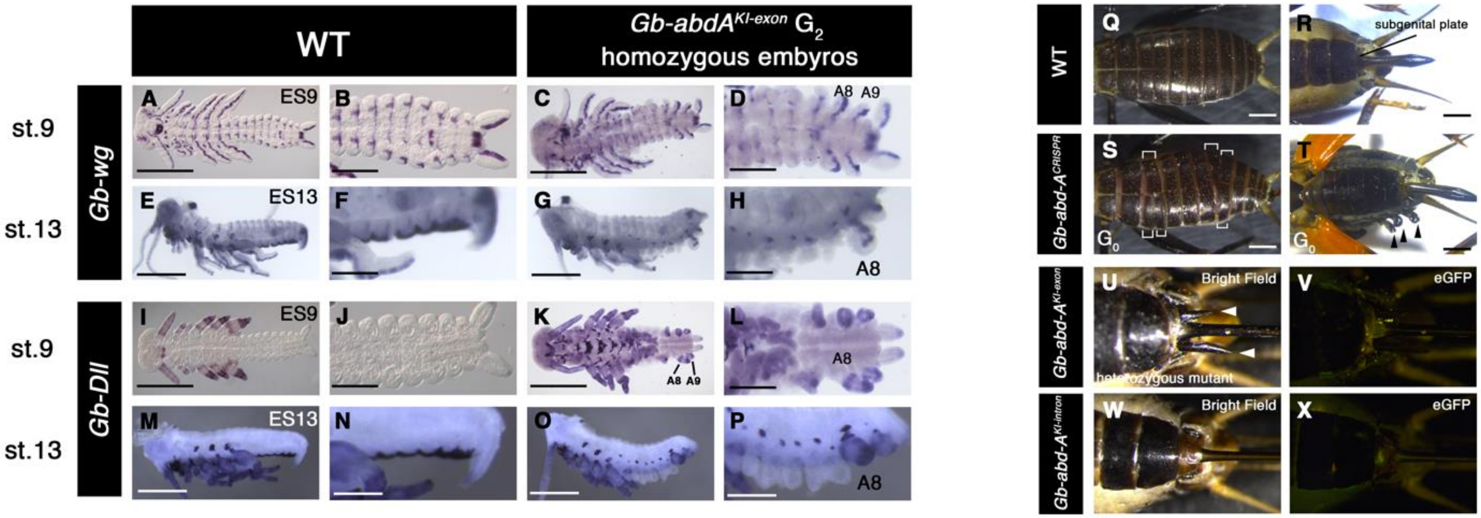
WT
Gb-abdA^KI-exon G2
homozygous embyros
st.9
st.13
Gb-wg
Gb-Dll
A ES9
B
C
D A8 A9
E ES13
F
G
H A8
I ES9
J
K A8 A9
L A8
M ES13
N
O
P A8
WT
Q
R subgenital plate
Gb-abd-A^CRISPR
S G0
T G0
Gb-abd-A^KI-exon
U Bright Field
heterozygous mutant
V eGFP
Gb-abd-A^KI-intron
W Bright Field
X eGFP

### Figure S7

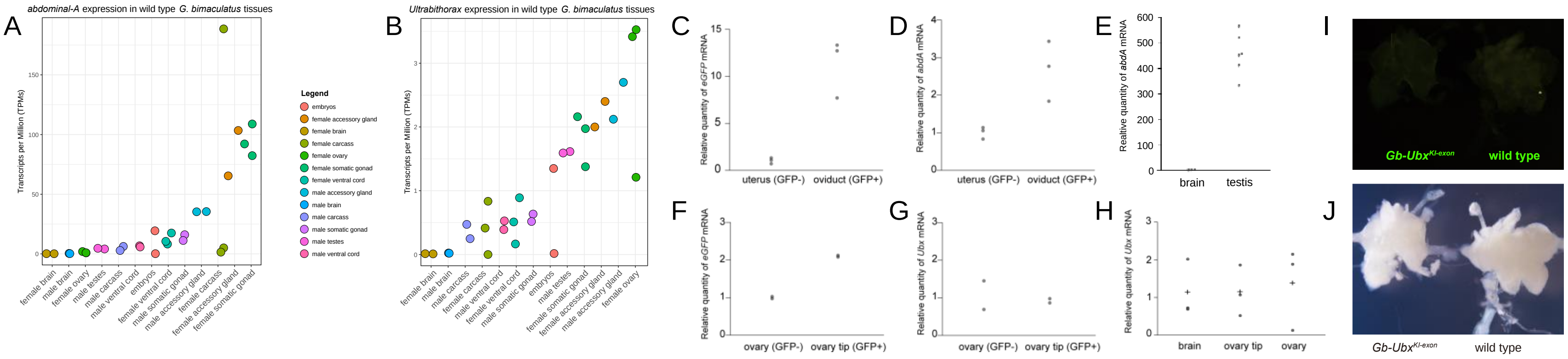
A
abdominal-A expression in wild type G. bimaculatus tissues
Transcripts per Million (TPMs)
B
Ultrabithorax expression in wild type G. bimaculatus tissues
Legend
embryos
female accessory gland
female brain
female carcass
female ovary
female somatic gonad
female ventral cord
male accessory gland
male brain
male carcass
male somatic gonad
male testes
male ventral cord
C
Relative quantity of eGFP mRNA
uterus (GFP-)
oviduct (GFP+)
D
Relative quantity of abdA mRNA
uterus (GFP-)
oviduct (GFP+)
E
Realtive quantity of abdA mRNA
brain
testis
I
Gb-Ubx KI-exon
wild type
F
Relative quantity of eGFP mRNA
ovary (GFP-)
ovary tip (GFP+)
G
Relative quantity of Ubx mRNA
ovary (GFP-)
ovary tip (GFP+)
H
Relative quantity of Ubx mRNA
brain
ovary tip
ovary
J
Gb-Ubx KI-exon
wild type

#### Figure S8

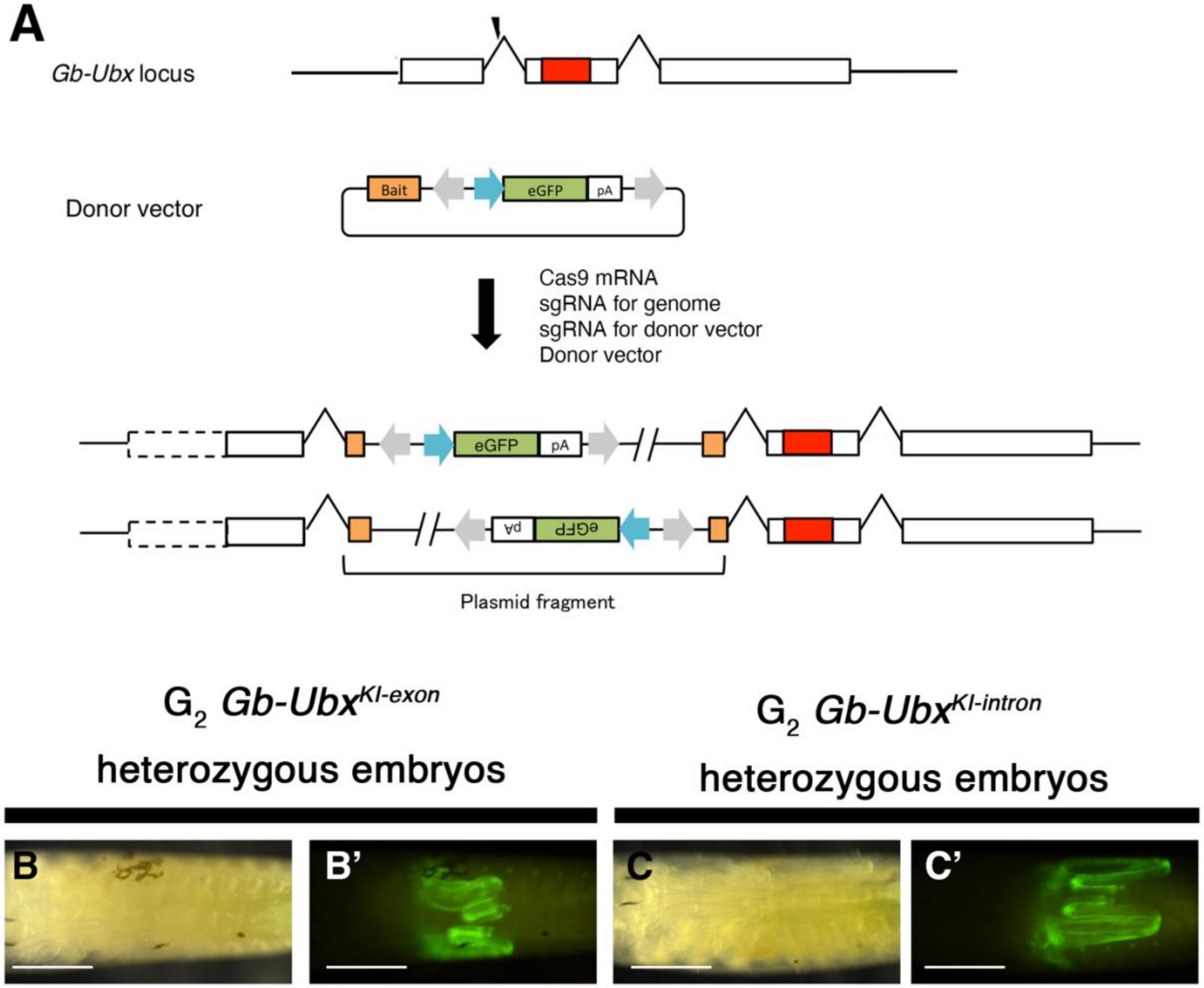
A
Gb-Ubx locus
Donor vector
Bait
eGFP
pA
Cas9 mRNA
sgRNA for genome
sgRNA for donor vector
Donor vector
Plasmid fragment
G2 Gb-Ubx KI-exon
heterozygous embryos
G2 Gb-Ubx KI-intron
heterozygous embryos
B
B'
C
C'
